# Dissetangling the Vine: Phylogenomics and Historical Biogeography of *Vanilla* (Orchidaceae)

**DOI:** 10.64898/2026.04.07.716943

**Authors:** Alexander Damián-Parizaca, Oscar Peréz-Escobar, Adam P. Karremans, Alexandre Antonelli, John P. Janovec, Nicole Mitidieri-Rivera, Olivia Jayne Fitzpatrick, Andres Barona, Xingbo Wu, Mathias Erich Engels, Marcelo Rodrigues Miranda, Wilfrido de la Cruz, German Carnevali, Gerardo Salazar, Eric Hágsater, Marilia C.R. Pappas, Daxs Coayla, Iván Tamayo-Cen, Rebeca Menchaca, Eric Smidt, Miguel A. Lozano-Rodriguez, Yader Ruiz, Leisberth Vélez, Henry Garzón, Luis Baquero, Gabriel Iturralde, Álvaro J. Pérez, Marco Jiménez, Segundo Oliva, Kenneth Cameron

**Affiliations:** Department of Botany, University of Wisconsin–Madison, 430 Lincoln Drive, Madison, Wisconsin 53706, U.S.A.; Inkaterra Asociación, Av. Víctor Larco Herrera 130, Miraflores, Lima, Peru; Royal Botanic Gardens, Kew, Richmond, Surrey, TW9 3AE, UK; Centro de Investigación Jardín Botánico Lankester, Universidad de Costa Rica, Cartago, Apartado 302-7050, Costa Rica; Gothenburg Global Biodiversity Centre, Department of Biological and Environmental Sciences, University of Gothenburg, Box 463, 405 30 Göteborg, Sweden; Department of Biology, University of Oxford, South Parks Road, Oxford, OX1 3RB, United Kingdom; Free State Growers, Inc. 12819 198^th^Street, Linwood, Kansas; Instituto Amazónico de Investigaciones Científicas SINCHI, Herbario Amazónico Colombiano Dairon Cárdenas López COAH, línea Flora-Sede Principal Leticia, Av. Vásquez Cobo entre calles 15 y 16, Leticia, Amazonas, Colombia; Horticultural Sciences Department, Tropical Research and Education Center, University of Florida IFAS, Homestead, FL, USA; Universidade Federal do Paraná, Departamento de Botânica, Programa de Pós-graduação em Botânica, Cx. P. 19031, Jardim das Américas, Curitiba, Paraná, Brazil; Companhia de Saneamento Básico do Estado de São Paulo, Caraguatatuba, SP, Brazil; Fundación Pachamama, Av. Alfonso Lamiña, El Potrero de San Luis de Lumbisí, Oficina 5, Quito, Ecuador; Centro de Investigacion Cientifica de Yucatan, Merida, Mexico; Departamento de Botánica, Instituto de Biología, Universidad Nacional Autónoma de México, Apartado Postal 70-367, 04510 México City, Mexico; Herbario AMO, Montañas Calizas 490, Lomas de Chapultepec, Miguel Hidalgo, CDMX, 11000, México; Embrapa Recursos Genéticos e Biotecnologia, Brasília – DF, Brazil; Centro de Investigaciones Tropicales-Universidad Veracruzana, México; Universidade Federal do Paraná, Setor de Ciências Biológicas, Departamento de Botânica, Laboratório de Sistemática e Ecologia Molecular de Plantas, Centro Politécnico, Caixa Postal 19031, Curitiba, PR, 81531–970, Brasil; Universidad de El Salvador, San Salvador, El Salvador; Herbario HUTPL, Departamento de Ciencias Biológicas, Universidad Técnica Particular de Loja, San Cayetano Alto s/n 11-01-608, Loja, Ecuador; Grupo de Investigación en Biodiversidad, Medio Ambiente y Salud (BIOMAS), Carrera de Ingeniería en Agroindustria, Facultad de Ingenierías y Ciencias Aplicadas, Universidad de Las Américas, UDLA, Vía a Nayón, Quito 170124, Ecuador; Herbario QCA, Escuela de Ciencias Biológicas, Pontificia Universidad Católica del Ecuador, Av. 12 de octubre 1076 y Roca, Apartado 17-01-2184, Quito, Ecuador; Universidad Nacional Toribio Rodriguez de Mendoza, Amazonas, Peru

**Keywords:** Angiosperm353, ILS, Neotropics, reticulate evolution, Vanilloideae

## Abstract

Renowned for its aromatic fruits and economic importance, the genus *Vanilla* poses longstanding taxonomic and phylogenetic challenges. Despite recent molecular studies, a comprehensive species tree is lacking, and the evolutionary processes and historical patterns shaping the genus remain poorly understood.

We present a new, comprehensive phylogenomic framework for *Vanilla*, based on 349 low-copy nuclear genes and 76 plastid loci from the Angiosperms353 probe set, which we used to infer evolutionary relationships, assess cyto-nuclear and gene–species tree discordance, and thoroughly investigate its historical distribution and diversification.

Sampling 43% of the genus, our framework resolves phylogenetic uncertainties, clarifies major clades, confirms prior hypotheses, and reveals novel placements, including *V. planifolia* and *Vanilla* subg. *Gondwana*. Discordances are primarily driven by incomplete lineage sorting, particularly in the vanillin-producing clade, with evidence of both ancient and recent hybridization, including a natural hybrid from the Yucatán Peninsula. Biogeographic analyses indicate a Guiana Shield origin (∼30 Mya), Amazonia as a major diversification source, the Andes as a permeable barrier, and Central America as the main diversification sink.

This study provides a robust evolutionary framework for *Vanilla*, supporting taxonomic revisions, comparative trait analyses, and a deeper understanding of the processes shaping this economically and biologically important orchid genus.

## Introduction

Vanilla is one of the most important natural flavorings from the Americas and remains one of the main orchid-derived spices recognized worldwide (Correll *et al.*, 1953). Naturally obtained from the fruits of the *Vanilla* Mill. plant and valued by Mesoamerican cultures since pre-Hispanic times (Prescott, 2005), Vanilla remains economically and culturally important, with a global organic market estimated at USD 1.26 billion (FAO, 2025). Although only *Vanilla planifolia* Andrews is widely cultivated as a crop, the genus comprises more than 100 species, pantropically distributed and particularly diverse in the Neotropics, where its highest diversity has been reported (Karremans *et al.*, 2020; Damian *et al.*, 2025; Fig. 1i). Representing the largest radiation within Vanilloideae and the most species-rich lineage among the deeper orchid subfamilies (Cameron, 2003, 2007; Chase *et al.*, 2015), members of *Vanilla* exhibit atypical traits for orchids, which have historically complicated its classification (Cameron, 2003). Although unique synapomorphies remain difficult to define, the genus is consistently recovered as monophyletic and characterized by diagnostic traits, notably a nomadic vine habit and berry-like fruits containing wingless, crustose black seeds (Fig. 1a–c; Soto Arenas, 2003; Soto Arenas & Cribb, 2010; Karremans *et al.*, 2025). Despite extensive historical research, many aspects of Vanilla biology and ecology remain poorly understood. At the same time, the genus faces increasing threats from habitat loss and climate change (Watteyn *et al.*, 2025), and its cultivation is limited by pest susceptibility, climate stress, and low genetic diversity resulting from clonal propagation (Flanagan *et al.*, 2018; Chambers *et al.*, 2021; Serna-Sanchez *et al.*, 2025). A robust understanding of evolutionary relationships within *Vanilla* is therefore critical to guide conservation, sustainable use, and the genetic improvement of the genus.

**Fig 1.**
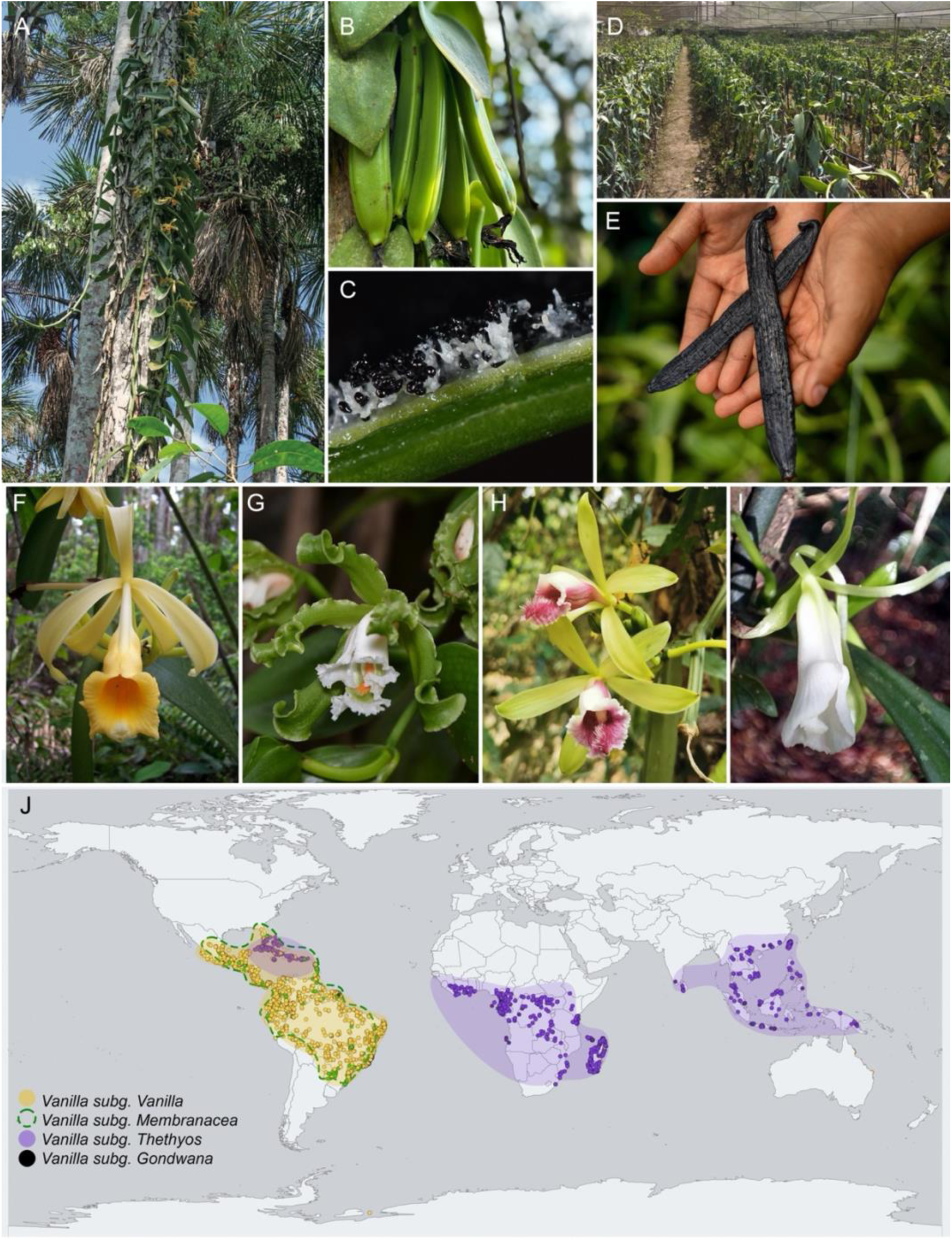
Morphology, distribution, and representatives of *Vanilla*. Morphological features: (A) Habit of *Vanilla pompona* inhabiting Amazonian wetlands, Peru; (B) Fruits of *V. pompona*; (C) Seeds of *V. armoriquensis*. Crop production of *Vanilla*: (D) Cultivation facility of *V. pompona*, Peru; (E) Curated fruits of *V. pompona*, Peru. Representatives of *Vanilla* subgenera: (F) *Vanilla* subg. *Vanilla* (*V. pompona*); (G) *Vanilla* subg. *Membranacea* (*V. andina*); (H) *Vanilla* subg. *Thethyos* (*V. cruenta*); (I) *Vanilla* subg. *Gondwana* (*V. cameroniana*). (J) Global distribution of *Vanilla* species, sourced from GBIF. Images: John Janovec (A), Alexander Damian (C), Arturo Rivas (B,D,E), Henry Garzon (G), Sukanya Thanombuddha (H), (I) Cyril Gaertner.

Efforts to resolve evolutionary relationships in *Vanilla* began in the late 1990s and early 2000s with several Sanger-based studies that clarified its placement within Orchidaceae and identified major clades (Cameron, 1996; Cameron & Chase, 1999, 2000; Cameron *et al.*, 1999; Soto Arenas, 2003; Cameron &Molina, 2006; Cameron, 2004–2009; Bouetard *et al.*, 2010). Building on this foundation, Soto Arenas &Cribb (2010), using rbcL sequences (Soto Arenas, 2003), proposed the first DNA-based infrageneric classification of *Vanilla*, recognizing three major clades distinguished primarily by leaf characters and a few floral traits: *Vanilla* subg. *Vanilla* and *Vanilla* subg. *Xanata* Soto Arenas and Cribb, the latter further divided into *Vanilla* sect. *Xanata* and *Vanilla* sect. *Thethya* Soto Arenas and Cribb. However, the recent proposal to conserve *V. planifolia* as the type of the genus (Karremans &Pupulin, 2023), together with the absence of well-defined synapomorphies for the inner major lineages, highlighted the need to revise the Soto Arenas and Cribb framework. In response, our research group integrated Sanger sequencing data from multiple DNA regions with an expanded morphological analysis, recognized four subgenera that more accurately reflect major evolutionary relationships (Karremans *et al.*, 2025): *Vanilla* subg. *Vanilla*, *Vanilla* subg. *Membranacea* Karremans, Pupulin and Damian, *Vanilla* subg. *Thethyos* Karremans, Damian and Pupulin, and *Vanilla* subg. *Gondwana* Karremans, Damian and Pupulin.

While these studies represent substantial progress toward a stable infrageneric classification, relationships at the species level remain largely unresolved. Most Sanger-based phylogenies remain characterized by persistent polytomies, low support values, and conflicting topologies among loci and analyses (Pansarin, 2016; Villanueva-Viramontes *et al.*, 2017; Ellestad *et al.*, 2022; Pansarin and Ferreira, 2022; Andriamihaja *et al.*, 2023; 2024; Karremans *et al.*, 2023; Cascales *et al.*, 2023; Krahl *et al.*, 2024). Consequently, the placement of several key taxa remains unstable, including the widely cultivated *V. planifolia*, whose position varies across single-locus and multilocus datasets, as well as numerous other species (Soto Arenas, 2003; Soto Arenas & Dressler, 2010; Bouetard *et al.*, 2010; Cameron, 2011a,b, 2019; Pansarin & Ferreira, 2022; Pansarin *et al.*, 2023). This lack of resolution is further exacerbated by uneven taxon sampling, particularly in Andean and African lineages (Damian *et al.*, 2025; NCBI, 2025). Additionally, chromosome-scale *Vanilla* genomes now provide powerful resources for studies of genetic diversity and crop improvement (Hasing *et al.*, 2020; Piet *et al.*, 2022) and hold clear promise for high-throughput phylogenomic inference (Hemstrom *et al.*, 2024). However, these tools have not yet been fully exploited to produce a comprehensive, well-supported, and reproducible evolutionary framework for the genus.

A major consequence of limited and uneven data sampling in *Vanilla* is pervasive phylogenetic incongruence, arising from both stochastic processes and biological factors that remain insufficiently assessed in the genus. Missing gene data, particularly when nonrandomly distributed, can bias inferred relationships and exacerbate stochastic error, with serious implications for phylogenetic inference (Wendel & Doyle, 1998; Lemmon *et al.*, 2009; Nabhan & Sarkar, 2012; Simmons, 2014; Steenwyk *et al.*, 2023;). Compounding these issues, many phylogenetic studies, including those focused on *Vanilla*, exhibit limited reproducibility owing to the frequent unavailability of raw data (e.g., DNA sequences, alignments, and inferred trees) and inadequate reporting of analytical parameters (e.g., software versions, substitution models, search strategies, and random seeds). Phylogenetic irreproducibility has itself been proposed as a contributor to phylogenomic incongruence, potentially influenced by variation in thread configurations, floating-point arithmetic behavior, and compiler differences (Shen *et al.*, 2020). Together with persistent challenges in taxonomic identification (Karremans et al. 2020), these factors substantially hinder efforts to reconstruct a robust and reliable phylogeny of *Vanilla* based on currently available datasets.

Phylogenetic incongruence may also arise from biological processes, such as ILS, hybridization, and introgression (e.g., chloroplast capture) (Madison, 1997; Smith *et al.*, 2015; Mirarab *et al.*, 2021; Molloy & Warnow, 2023; Steenwyk *et al.*, 2023). *Vanilla* is well-suited to investigating these processes, as previous phylogenetic inferences have consistently depicted relationships within *Vanilla* as characterized by short internodes and recent divergence times (e.g., Bouétard *et al.*, 2010; Villanueva-Viramontes *et al.*, 2017), both of which are well-established predictors of elevated ILS (Degnan and Rosenberg, 2006). Similarly, the genus exhibits limited genetic barriers, widespread natural and cultivated interspecific hybridization (Rodolphe *et al.*, 2011; Ramos-Castella *et al.*, 2016; Hu *et al.*, 2019; Pansarin, 2025), and successful introgression of both male and female traits in artificial crosses (Divakaran *et al.*, 2006). Together, these characteristics suggest that gene tree discordance in *Vanilla* likely reflects a complex interplay of ILS and reticulate evolution, rather than solely methodological artifacts.

Another area of scientific inquiry into *Vanilla* that is impeded by phylogenetic incongruence and a lack of a well-resolved phylogeny is historical biogeography. Early pre-DNA hypotheses placed the center of *Vanilla* diversification in Indo-Malaysia, followed by multiple dispersal events to the Pacific, the Americas, and Africa (Portères, 1951, 1954), but molecular phylogenetic studies later suggested an American origin with early transcontinental migration predating the breakup of Gondwana (Cameron, 1999, 2000). Fossil-calibrated analyses dated the crown node of *Vanilla* to ∼34.5 Ma, effectively rejecting Gondwanan vicariance and supporting a Neotropical origin (Ramírez *et al.*, 2007; Bouetard *et al.*, 2010). Under this scenario, the pantropical distribution of the genus is explained by three major transoceanic dispersal events: from the Neotropics to Africa (∼25 Ma), from Africa to Asia (∼10.2 Ma), and from Africa back to the Neotropics (∼10.8 Ma), potentially mediated by floating vegetation, wind, and avian vectors across oceanic corridors such as the Rio Grande Rise–Walvis Ridge and the Northern Atlantic Land Bridge. While these broad patterns are supported by plastid-based analyses (Givnish *et al.*, 2016), more recent integrative studies combining Sanger and phylogenomic data propose a Laurasian–Neotropical origin for Vanilloideae and a Nearctic–Neotropical origin for *Vanilla*, with a younger crown age of ∼30 Ma (Pérez-Escobar *et al.*, 2024). However, all of these studies remain limited by sparse species-level sampling, unidentified or misidentified accessions, and in some cases inferred or grafted relationships, restricting the precision of current biogeographic reconstructions (Givnish *et al.*, 2016; Karremans *et al.*, 2020; Pérez-Escobar *et al.*, 2024).

Although previous studies have advanced our understanding of *Vanilla* evolution, the primary biogeographical drivers shaping its distribution remain poorly understood, particularly within internal lineages where phylogenetic uncertainty persists. For example, the Neotropics, often regarded as the center of origin for the genus, lacks up-to-date, explicit biogeographic analyses, and limited taxon sampling continues to hinder robust interpretations of its distributional dynamics. For instance, *Vanilla* subgenus *Membranacea*, which comprises approximately 36% of Neotropical *Vanilla* diversity, has been largely neglected in phylogenetic studies (Damian *et al.*, 2025), with most analyses including only a single accession (*V. mexicana* Mill.), leaving both the crown age and evolutionary history unresolved. Preliminary phylogenetic hypotheses with denser sampling suggest at least three distinct subclades within *Membranacea*, each exhibiting divergent distribution patterns, indicative of a complex biogeographic history (Damian *et al.*, 2024; Karremans *et al.*, 2025). Similar gaps exist in other species-rich regions, such as the Indian Ocean and continental Africa (Andriamihaja *et al.*, 2020), where orchid lineages with *Vanilla*-like distributions display intricate biogeographical patterns (Freudenstein, 2024). Expanded taxon sampling may also uncover additional long-distance dispersal events, as recently proposed for several leafless *Vanilla* species from Madagascar (Andriamihaja *et al.*, 2023). Likewise, unsampled or phylogenetically unplaced taxa, such as the leafless Neotropical *V. penicillata* Garay and Dunst. or the recently described *V. cameroniana* Damian, with their distinctive traits, hold potential not only to clarify evolutionary relationships within *Vanilla* but also to illuminate the genus’s complex biogeographic history.

Here, we present the most comprehensive phylogenomic framework of *Vanilla* to date. Through a broad collaborative effort, we maximized taxon sampling, including 43% of the currently recognized species diversity. Our objectives were as follows: (1) reconstruct a robust phylogeny of *Vanilla* using global representative sampling with the Angiosperm353 probe set and inferring nuclear and plastid multigene species trees under both coalescent-based and supermatrix approaches; (2) quantify phylogenetic discordance and evaluate the roles of ILS and hybridization within the genus; and (3) infer the biogeographic history of *Vanilla*. This study will provide a robust evolutionary framework for future systematic and evolutionary research on *Vanilla*, facilitating deeper exploration of its remarkable biological and economic significance.

## MATERIALS AND METHODS

### Taxon sampling

Overall, we were able to sample 204 taxa representing 54 unique *Vanilla* species in all subgenera *Membranacea, Vanilla, Gondwana* and *Thethyos*. In most cases, we included multiple accessions from different countries to assess species boundaries, averaging five per taxon and up to eleven in the case of *Vanilla pompona* Schiede. We complemented our ingroup sampling with seven *Vanilla* accessions from a previous broad orchid phylogenomic study which used the Angiosperm353 (Perez-Escobar *et al.*, 2024). As outgroups we included representatives of every accepted genus within both Pogonieae and Vanilleae tribes in Vanilloideae (with the exception of *Eriaxis* Rchb. f.), as well as three species within Cypripedioideae (*Phragmipedium lindleyanum* (R.H. Schomb. ex Lindl.) Rolfe*, P. warszewiczianum* (Rchb. f.) Schltr., *Selenipedium aequinoctiale* Garay) and one within Apostasioideae (*Apostasia wallichii* R. Br.), these mainly from data generated for this study and complemented by previous work (Perez-Escobar *et al.*, 2024). All complemented data were downloaded manually from NCBI (https://www.ncbi.nlm.nih.gov/) and ENA (https://www.ebi.ac.uk).

Samples for this study were obtained from four sources. First, 146 were extracted from fresh silicagel-dessicated tissue samples collected exclusively for this study. Another 22 taxa from herbarium material, an additional 32 taxa were represented by DNA aliquots generated by Cameron (1996, 2009). Four taxon samples were provided by Miguel Angel Soto Arenas (UNAM). In addition, we were able to sample and sequence type material from six taxa (*V. andina* Damián and H. Garzón, *V. armoriquensis* Damian and Mitidieri*, V. cameroniana, V. cribbiana* Soto Arenas, *V. javieri inedit*., and *V. sekut* Damián, H. Garzón and A. Bentley). All voucher and source details are provided in Appendix S1.

### DNA extraction, library preparation and sequencing

DNA was extracted from between 40 and 100 mg of silica-dried or herbarium tissue, depending on the availability of starting tissue. Dried samples were powdered in a Retsch MM 400 shaker for five minutes and extracted using a modified CTAB extraction method (Doyle and Doyle, 1987). Extractions were resuspended in 70µl of 10 mM Tris buffer. The resulting DNA was quantified using a Quantus fluorometer (Promega, Madison, WI, USA) and fragment size was assessed on a 1.5% agarose gel stained with SYBR Safe. Samples were diluted to 200 ng in 10 mM Tris. With the exception of some herbarium samples, DNA fragments were exceeded 350 bp and were therefore sonicated using a Covaris ME220 Focused Ultrasonicator (Covaris, Woburn, MA, USA) with Covaris microTUBES to obtain a fragment size range of 350 - 450 bp. Fragment size distribution were subsequently verified using a TapeStation 4200 system (Agilent Technologies, Santa Clara, CA, USA) with D1000 tapes.

Libraries were prepared using the NEBNext Ultra II DNA Library Prep Kit (New England BioLabs, Ipswich, MA, USA). Size selection was omitted for samples with highly degraded DNA such as those derived from herbarium material. Libraries were prepared following the manufacturers protocol, with reaction volumes reduced by half throughout. DNA extractions and libraries were prepared at the Cameron lab (University of Wisconsin-Madison), Kew Gardens, the Laboratório de Sistemática e Ecologia Molecular de Plantas (Universidade Federal do Paraná), and Instituto de Biología (Universidad Nacional Autónoma de Mexico). Subsequent quality control, hybridization, and sequencing using the Angiosperms353 probe set (Johnson et al. 2018) were conducted at Daicel Arbor Biosciences following standard protocols.

### Data processing and loci filtering

We assembled two major datasets. For the nuclear dataset, paired–end DNA raw reads were adapter–trimmed and quality–filtered using TrimGalore v.0.6.4 (available at (https://github.com/FelixKrueger/TrimGalore), applying a Phred score quality threshold of 30, a minimum read length of 20, and retaining only read pairs that passed all quality–filtering criteria. For each sample, Angiosperms353 loci were retrieved using the HybPiper v.2.3.3 pipeline (Johnson *et al.*, 2016). To improve the recovery of nuclear marker, we employed the taxonomically extended Mega353 target file (McLay *et al.*, 2021). Clean reads were mapped using both Burrows–Wheeler Alignment (BWA) v.0.7 (Li & Durbin, 2009) and Diamond (Buchfink *et al.*, 2015). Although the latter approach was more time- and computationally intensive, it recovered a greater number of loci and yielded longer average sequence lengths; therefore, Diamond-derived outputs were used for all downstream analyses.

Mapped reads were assembled de novo for each gene separately using SPAdes v. 3.13 (Bankevich *et al.*, 2012), with a minimum coverage threshold of 6×. Retrieved sequences were subsequently filtered by length using the *filter_by_lenght* flag in HybPiper with a – percent_filter value of 40%. Putative paralogs were identified and flagged using the HybPiper-built paralog_retriever module and excluded using a custom script. Genes represented by fewer than five taxa were removed, as were taxa containing less than 10% of the total set of 353 target genes. For each locus, homologous sequences were combined and aligned using MAFFT v. 7.4 (Katoh & Standley, 2013) under the FFT–NS– i strategy. Gene alignments were subsequently trimmed to remove spurious sequences and excessive missing data using trimAL v.1.2 (Capella–Gutiérrez *et al.*, 2009) with the *-automated1* option.

For the Plastome dataset, we employed the CAPTUS v1.3.3. pipeline (Ortiz *et al.*, 2023), which, in combination with the non-specific universal Angiosperm353 probe set, enabled the recovery of a substantial number of off-target reads. Raw sequence data were subjected to adapter trimming and quality filtering using BBTools (available at sourceforge.net/projects/bbmap) under default parameters. All adapter sequences were removed, leading and trailing bases with PHRED scores below 13 were trimmed, and reads with an average PHRED score below 16 were discarded. Clean reads were assembled using MEGAHIT v1.2.9 (Li et al. 2015) with default CAPTUS (CAPSKIM) settings. Plastid genes were then extracted from the assembled contigs using the universal seed plant plastid protein-coding reference set implemented in CAPTUS (*--ptd_refs SeedPlantsPTD*). Final gene alignments were generated using MAFFT with default parameters, including paralog-filtering procedures. Aligned but unfiltered sequences were subsequently quality-trimmed using ClipIT v.2.1.1. (Steenwyk *et al.*, 2020), which removes empty and excessively short alignment columns.

### Phylogenetic analyses

Phylogenetic relationships were reconstructed using both coalescent-based species tree and concatenation approaches for the Nuclear dataset, whereas only a concatenation approach was applied to the nonrecombinant Plastome dataset. For the coalescent analysis, maximum likelihood (ML) gene trees were inferred for each nuclear locus with IQ-TREE v3.0.1 (Wong *et al.*, 2025). The best-fit substitution model for each locus was selected with ModelFinder (Kalyaanamoorthy et al. 2017) under the Bayesian Information Criterion (BIC). Branch support was evaluated using 1,000 ultrafast bootstrap replicates (Hoang *et al.*, 2018).

To account for topological incongruence among nuclear gene trees, a species tree was estimated under the multispecies coalescent (MSC) model in ASTRAL-IV, as implemented in ASTER (Zhang *et al.*, 2025). Gene tree branches with likelihood bootstrap support (LBS) values below 20 were collapsed using Newick Utilities (Junier & Zdobnov, 2010), following Sayyari & Mirarab (2016). The final ASTRAL tree was annotated with detailed branch support (option -u 2) and rooted using *Apostasia wallichii*.

Concatenated matrices for the Nuclear and Plastome datasets were generated using the *concat* function of the Python-based program AMAS (Borowiec, 2016), with partition information retained. Maximum likelihood analyses of the concatenated datasets were conducted in IQ-TREE using the *-p* option to specify partitions and *-m* MFP+MERGE to optimize model selection and partitioning schemes. Branch support was assessed with 1,000 ultrafast bootstrap replicates. All ML analyses were performed on the Center for High Throughput Computing (CHTC) servers at the University of Wisconsin–Madison.

### Genetic distinctness

To complement the phylogenetic analyses, overall sequence divergence was assessed using multidimensional scaling (MDS) to explore patterns of differentiation among taxa. Pairwise genetic distances were computed under the Kimura 2-parameter model (Kimura, 1980) using the dist.dna function in the ape package (Paradis & Schliep, 2019) in R, based on concatenated post-trimmed alignments with pairwise deletion enabled. The two axes explaining the greatest proportion of variance were retained for visualization. An initial MDS plot including all accessions was generated to identify major groupings, followed by a second analysis restricted to *Vanilla* taxa, in which data points were labeled by subgenus. Both plots were produced in RStudio using ggplot2 (Wickham, 2016). As noted by de Vos *et al*. (2025), MDS is a phenetic approach that provides complementary evidence for assessing taxon differentiation by summarizing the structure of sequence variation and facilitating interpretation of phylogenetic patterns.

### Phylogenomic and gene-tree discordance

Intergenomic discordance between the Nuclear and Plastid datasets was visualized using phylogenetic tanglegrams generated with the cophylo function in *phytools* (Revell, 2024). Comparisons were conducted between the Nuclear ASTRAL species tree and the Plastome ML tree, as well as between the Nuclear ML and Plastome ML and both ML Nuclear and Plastome trees. Genealogical concordance across nuclear loci was quantified using gene and site concordance factors (gCF and sCF) implemented in IQ-TREE v3.0.1, employing the ASTRAL species tree, post-trimmed gene alignments, and the corresponding ML gene trees as inputs. In addition, polytomy tests were performed in ASTRAL-III (Zhang *et al.*, 2018) using the *-t 10* option. Under this framework, the null hypothesis posits that a given branch in the species tree represents a polytomy given the set of gene trees; a p-value > 0.05 indicates failure to reject the null hypothesis, consistent with the presence of polytomies while accounting ILS. Results were visualized in R using ggplot2.

Finally, phylogenetic conflict was further assessed in SplitsTree v6.5.1 (Huson & Bryant, 2006) by constructing a NeighborNet network based on uncorrected *p*-distances derived from the concatenated nuclear alignments.

### Hybridization and ILS detection

We used *PhyTop* v1.0 (Shang *et al.*, 2025) to visualize quartet scores (*q₁, q₂, q₃*) derived from the ASTRAL analyses and to detect signals of introgressive hybridization (IH) and ILS. *PhyTop* tests the null hypothesis that the frequencies of the two discordant quartets are equal (*q₂ = q₃*), which is indicative of ILS, and computes indices quantifying the strength of ILS and IH (ILSi, IHi) as well as the proportion of quartet variation explained by each process (ILSe, IHe). Nodes with ILSi < 0.3 and IHi < 0.05 were considered highly supported. Nodes with Ihi > 0.2 and p < 0.05 were interpreted as showing IH, whereas nodes with ILSi > 0.7 and p > 0.05 were interpreted as exhibiting strong ILS signals.

To further test for IH, we employed *PhyloNet* v3.8.2 (Wen *et al.*, 2018) using a reduced taxon set comprising clades previously identified as showing IH signals. Gene trees were subset with *Phyx* (Brown *et al.*, 2017), with the *-pxrmt* option limiting each analysis to a maximum of 14 taxa, and were rooted with *Apostasia wallichii* using the *-pxrr* option; trees lacking the outgroup were excluded. Phylogenetic networks were inferred under the maximum pseudo-likelihood model (InferNetwork_MPL) with ten replicates for each reticulation level (H₀–H₆), returning the five best networks per level. The optimal network was selected using a slope-heuristic approach (Cao *et al.*, 2019; Kong *et al.*, 2025).

In addition, we used *HyDe* v1.0.2 (Blischak *et al.*, 2018) to detect potential introgression within *Vanilla* and its allies by conducting exhaustive tests among all phylogenetic terminals based on the Nuclear dataset. Analyses were implemented using the Python script run_hyde.py with the *-ignore_amb_sites* flag enabled. Tests with significant *p*-values after Bonferroni correction (1 < γ > 0, *Z*-score > 3) were considered strong evidence of hybridization. We then visualized the frequency of terminals inferred to have hybrid origins and the distribution of γ-values using violin plots across multiple taxonomic levels (tribe, genus, infrageneric groups within *Vanilla*, and species). Additionally, we generated histograms summarizing the overall γ distribution for the entire dataset and for each terminal inferred to be of hybrid origin. Finally, we used *VisualHyDe* (available at https://github.com/Jhe1004/VisualHyDe) to visualize and interpret the detected gene flow events, providing an intuitive and comprehensive overview of hybridization patterns across taxa.

### Divergence Time Estimation

To infer a Bayesian time-calibrated phylogeny, we employed a gene-shopping approach by subsampling the nuclear dataset to identify the most clock-like loci using SortaDate (Smith *et al.*, 2018). Total tree length, root-to-tip variance, and topological congruence with the species tree were used as key criteria for assessing phylogenetic informativeness and the stability of molecular rates. The 20 highest-ranking genes were selected for downstream dating analyses to ensure a computationally tractability in Bayesian inference. The species tree recovered from ASTRAL analyses was used as the reference topology. Prior to running SortaDate, genes were rooted using the *pxrr* script within the *Phyx* package, with *Apostasia wallichii* designated as the outgroup; only genes successfully rooted with this outgroup were retained. Alignments for the top 20 genes were concatenated using the concat function in AMAS and analyzed with ModelFinder in IQ-TREE as a single partition to determine the best-fitting substitution model for subsequent BEAST analyses.

We calibrated the nuclear phylogeny using BEAST v.10.5.0 (Suchard *et al.*, 2018), employing a single partition and the GTR+F+I+R5 substitution model, with branch rates estimated under an uncorrelated relaxed clock and a birth–death tree prior. The posterior distribution of trees was constrained to include the nodes of the ASTRAL topology. Nine age priors, primarily based on Givnish *et al*. (2016), were applied as normally distributed priors with a standard deviation of 1 (Table S3), with the *autoOptimize* option enabled and the tree topology fixed. The analysis was run twice, totaling 300 million MCMC steps, with trees and parameters logged every 1,000 generations. Log and tree files were subsequently combined using LogCombiner v.10.5.0. Convergence of parameter estimates was assessed in Tracer v.1.7.2, and the Maximum Clade Credibility (MCC) tree was generated in TreeAnnotator v.10.5.0 with a 25% burn-in.

### Historical Biogeography

To investigate the historical biogeography of *Vanilla* and its close relatives, we compiled presence–absence data for each species across all recognized biogeographic regions. Distributional information was assembled primarily from herbarium specimens accessed through the Global Biodiversity Information Facility (GBIF) and the World Checklist of Vascular Plants (POWO, 2026). All outlier records were scrutinized individually, and specimen images were examined to verify taxonomic identifications; records lacking reproductive structures were removed due to the heightened risk of misidentification. Supplementary occurrence data were obtained from the published literature, with Karremans *et al*. (2020, 2025) and Damián *et al*. (2025a, 2025b) providing particularly critical insights for refining and corroborating species-level geographic distributions.

Ancestral area reconstruction was performed using the BEAST time-calibrated phylogeny, with a single accession per species. *Cleistesiopsis divaricata* and *Vanilla cameroniana* were grafted onto this tree according to their phylogenetic positions inferred from the Plastome tree. Similarly, *Vanilla africana*, *V. organensis*, and *V. verrucosa* were incorporated into the BEAST phylogeny based on positions inferred by Bouetard et al. (2010) and Krahl et al. (2025). Analyses were conducted in the R package BiogeoBEARS (Matzke, 2013), which allows the simultaneous evaluation of alternative biogeographic models to assess the relative contributions of dispersal, local extinction, vicariance, founder-event speciation, and within-area speciation to present-day distributional patterns. Three models were evaluated independently: (1) DEC (dispersal–extinction–cladogenesis; Ree and Smith, 2008), which estimates dispersal and extinction rates and models range inheritance at speciation events under a likelihood framework; (2) DIVA-Like (modified from Ronquist’s 1997 dispersal–vicariance analysis), which permits dispersal and extinction along branches and vicariant range subdivision at cladogenesis; and (3) BayArea-like (Landis *et al.*, 2013), a likelihood-based framework designed for datasets with numerous geographic areas that characterizes range evolution primarily as a function of within-lineage processes. For each model, paired analyses were conducted with and without the founder-event speciation parameter (J), resulting in six total model configurations. Model fit was compared using the corrected Akaike Information Criterion (AICc), and the best-supported model was identified based on the highest AICc weight.

To evaluate complementary biogeographic hypotheses while minimizing model overparameterization, two hierarchical sets of biogeographic regions were defined. The first set, applied to the full vanilloid dataset, comprised six broad regions intended to capture large-scale, transcontinental dispersal patterns: Nearctic (A), Neotropics (B), Palearctic (C), Afrotropics (D), Australasia (E), and Indo-Malaya (F). The second set, restricted to *Vanilla*, included eight regions providing finer spatial resolution within the predominantly Neotropical distribution of the genus: Afrotropics (A), Indo-Malaya (B), Antilles (C), Central America + Florida (D), Andes + Pacific (E), Amazonia (F), Dry Diagonal (G), and Atlantic Rainforest (H). These regional delimitations follow established comparable distributions (e.g., Givnish *et al.*, 2016; Thode *et al.*, 2019; Carvalho *et al.*, 2020).

Two sets of parameters were analyzed under each of the three biogeographic models (DEC, DIVALIKE, BAYAREALIKE): (1) a time-stratified analysis incorporating dispersal-rate scalars that reflect changes in the paleogeography of the Americas through time, applied exclusively to the *Vanilla*-only dataset, and (2) an unconstrained analysis without time stratification or region-specific dispersal scalars, applied to both the vanilloid-wide and *Vanilla*-only datasets. Only ancestral states permitted by the dispersal matrix were allowed. The evolutionary history of the group was divided into five temporal bins following Robertson (2009), Hoorn *et al*. (2010), and Jaramillo and Oviedo (2017): 0–2.5 Ma (modern geography), 2.5–7 Ma (pre-closure of the Isthmus of Panama), 7–10 Ma (Acre System), 10–23 Ma (Pebas System and low-elevation Andes), and 23–30 Ma (exposed and unexposed rocks of the Antillean arc). Dispersal-multiplier matrices followed Vásquez-Restrepo *et al*. (2024), assigning values ranging from ∼0 to 0.75 assigned within each temporal bin: 0.75 for dispersal between adjacent regions, 0.5 for non-adjacent regions, 0.25 for dispersal across major barriers (e.g., oceanic barriers, the Andes, or the Acre or Pebas systems), and ∼0 for inaccessible regions. Full details of the dispersal matrices are provided in Table S4-S6.

Finally, to identify which areas acted as sources and sinks of *Vanilla* diversification, we performed a time-stratified stochastic mapping analysis (Dupin et al. 2017) on the *Vanilla*-only BEAST tree. Fifty stochastic maps were used to estimate the average numbers of anagenetic range expansions, extinctions, range switches, cladogenetic expansions, and jump-dispersal events. We then calculated the total number of dispersal events, encompassing both anagenetic (branch-level) and cladogenetic (node-level) events, with event tables first enriched with inferred source areas. Dispersal events were extracted, distinguishing their origin (“sources”) and destination (“sinks”) areas, further divided into 5 Myr time bins, and visualized in bar plots for analysis. These data were also used to construct a biotic interchange network for improved visualization of dispersal patterns.

## RESULTS

### The first phylogenomic framework dataset for Vanilla

We generated genome-scale data for 204 taxa including 54 species of *Vanilla* and an additional 30 species of *Vanilla* relatives. Subgeneric representation within *Vanilla* included 60% (12/20) of the species of subg. *Membranacea*, 80.5% (29/36) of subg. *Vanilla*, 100% (1/1) of subg. *Gondwana*, and 17.4% (12/69) of subg. *Thetyos*. Among the *Vanilla* species sampled, 90% (49 species) had not been included in previous phylogenomic analyses (e.g., Pérez-Escobar *et al.*, 2024), and 20 were newly sequenced for this study. From these data, we assembled two datasets: (1) Nuclear dataset, based on 349 nuclear low-copy genes, and (2) Plastome dataset: a concatenated matrix of 76 plastid coding genes. For the Nuclear dataset, the mean number of raw reads per *Vanilla* sample was 94,031,020, ranging from 32,874,048 in *V. planifolia* to 396,678,890 in *V. ribeiroi* Hoehne. On average, 306 of the 353 targeted nuclear loci from the Angiosperms353 bait set were recovered per sample (range: 120 in *V. labellopapillata_125* to 343 in *V. poitaei_131*) (Table S1). After filtering, the final Nuclear_MSC dataset included 162 accessions and 349 loci (98% of the target set). The number of high-quality loci recovered per species ranged from 35 (*V. yanesha_234*) to 324 (*V. odorata_118*). Individual locus alignments ranged from 93–3,411 bp (mean: 689.9 bp) with missing data between 0–54% (mean: 13%). For the Plastome dataset, the mean alignment length per accession was 797.7 bp, ranging from

268.2 bp in *V. pompona_238* to 952.4 bp in *V. planifolia_106*. Of the 80 targeted plastid genes, an average of 68 were recovered per accession, with a maximum of 73 in *V. parvifolia_150*, *V. armoriquensis_122*, *V. appendiculata_228*, *V. sprucei_233*, and *V. cristato-callosa_141*, and a minimum of 40 in *V. pompona_238* (Table S2). The final post-filtering Plastome dataset comprised 150 accessions and 76 plastid genes. *Vanilla parvifolia_150* had the largest number of loci recovered (73) and *V. pompona_238* the fewest (29). Individual plastid locus alignments ranged from 87–7,476 bp (mean: 878.6 bp), with missing data from 0–59.5% (mean: 9.6%). All newly generated reads are available in NCBI BioProject [TBD], and all individual Nuclear and Plastome locus alignments are deposited in Dryad [TBD].

The Nuclear_MSC produced a fully resolved phylogeny of *Vanilla* and its allies (Figs. 2b, S1-2). At the major scale, *Vanilloideae* and its two tribes are resolved as monophyletic (LPP = 1), with the vast majority of backbone nodes highly supported (LPP = 1). The sole exception was the position of *Lecanorchis*, which was moderately support as sister to the rest of the *Vanilleae* tribe (LPP = 0.74). At the genus level, all groups were recovered as monophyletic and highly supported (LPP = 1), similar to the internal nodes depicting the relationships across the different genera (LPP = 0.89–1). Within *Vanilla* we identified three strongly supported major clades (LPP = 1; Fig. 2b) recognized previously as subgenera *Vanilla* subg. *Membranacea*, *Vanilla* subg. *Thethyos*, and *Vanilla* subg. *Vanilla*. This clustering was consistently recovered in the network analysis conducted using SplitsTree (Fig. 2a) and corroborated by the genetic distinctness analysis (Fig. S3). Moreover, we recovered all sampled *Vanilla* species as monophyletic and highly supported (LPP = 1), except for *Vanilla odorata*, which had low support (LPP = 0.4). Among *Vanilla* subgen. *Membranacea*, we identified three highly supported main subclades: one primarily endemic to the Atlantic Rainforest (Clade A; LPP = 1), another largely restricted to the western and eastern Andean slopes (Clade B; LPP = 1), and a third widely distributed across the Neotropics (Clade C; LPP = 1). Most internal relationships within the *Membranacea* clade are highly supported (LPP = 0.89–1), with few exceptions, including the clade composed of *V. dietschiana* Edwall and *V. parvifolia* Barb. Rodr. and the one comprising *Vanilla*_spnov_212, *V. acuta* Rolfe, *V. martinezii* Soto Arenas, and *V. mexicana*, both only moderately supported (LPP = 0.63). We observed a similar pattern in the *Thethyos* clade, in which most relationships were strongly supported (LPP = 1). Within this clade, we recovered two of the three sections of the subgenus recently proposed by our research group (Karremans *et al.*, 2025), both of which were monophyletic and highly supported (LPP = 1). The only relationship with low support involved the clade comprising the Antillean *Vanilla poitaei* Rchb. f. and *V. claviculata* Sw., together with the remaining members of *Vanilla* sect. *Aphylla* (LPP = 0.27).

**Fig. 2.**
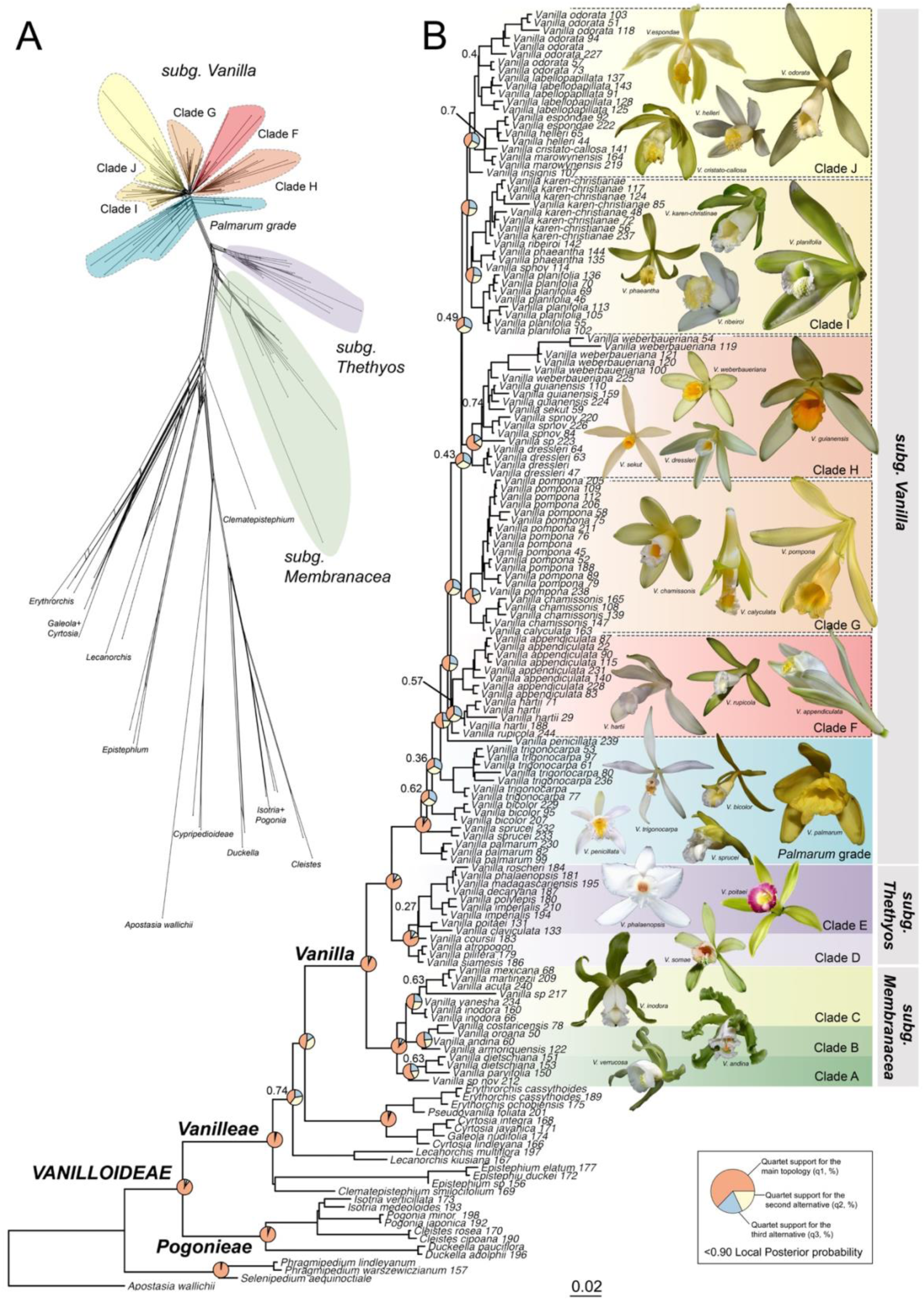
Phylogenetic network and species tree of *Vanilla*. (A) NeighborNet unrooted network including all genera of Vanilloideae. (B) Coalescent-based species tree of *Vanilla* estimated using ASTRAL-IV, showing quartet score pie charts on major backbone nodes. Local posterior probabilities (LPP) are indicated for nodes with values below 0.9. Different coloured blocks represents major clades.

Within the vanillin-producing clade (*Vanilla* subg. *Vanilla*), we identified five well-supported subclades and a basal grade (Fig. 1b), the latter forming a paraphyletic assemblage designated here as the “Palmarum grade.” This group includes species with strongly supported stem nodes for *V. penicillata*, *V. trigonocarpa* Hoehne, and *V. palmarum* (LPP = 1), whereas support for *V. sprucei* Rolfe and *V. bicolor* Lindl. is lower (LPP = 0.62 and 0.36, respectively). The remaining major subclades are all well supported (LPP = 0.92–1). At the deeper nodes, Clade F includes *V. appendiculata* Rolfe, *V. hartii*, and *V. rupicola* Pansarin and E.L.F. Menezes, with moderate support for the relationship between the first two species (LPP = 0.57). The next clade corresponds to Clade G, comprising *V. pompona*, *V. calyculata* Schltr., and *V. chamissonis* Klotzsch, with all internal relationships highly supported (LPP = 1). Following this, the third clade is recognized as Clade H, encompassing *V. cribbiana*, *V. dressleri* Soto Arenas, *V. sekut*, *V. guianensis* Splitg., *V. weberbaueriana* Kraenzl., and *V. janovecii*. Support within this section is heterogeneous with *V. dressleri* and *V. sekut* strongly supported (LPP = 1), whereas *V. janovecii.*, *V. guianensis*, and *V. weberbaueriana* exhibit short branches and low support (LPP = 0.25–0.35). Finally, two terminal subclades, Clades I and J, were recovered. Clade J includes *V. espondae* Soto Arenas, *V. helleri* A.D. Hawkes, *V. cristato-callosa* Hoehne, *V. marowynensis* Pulle, *V. labellopapillata* A.K. Koch, Fraga, J.U. Santos and Ilk.-Borg., *V. odorata* C. Presl, and *V. insignis* Ames, all strongly supported save for the relationship between *V. marowynensis* and the clade comprising *V. espondae* + *V. helleri* (LPP = 0.7). Clade I encompasses *V. planifolia* and its close relatives (*V. phaeantha* Rchb. f., *V. karen-christianae* Karremans and P. Lehm., *V. ribeiroi*, and *Vanilla_spnov_114*), with high support of all relationships (LPP = 0.8–1).

### Discordance within and among genomes and possible hybridization within Vanilla and its allies

Overall, both the Nuclear_ML and Nuclear_Astral analyses recovered largely congruent topologies, with only minor differences concentrated within the tribe Vanilleae. Within the Old World vanilloids, the placement of *Clematepistephium* N. Hallé differed between analyses: in the Nuclear_ML tree it was recovered as sister to the remainder of the tribe (BBS = 71), excluding *Epistephium* Kunth, whereas in the Nuclear_Astral analysis it was recovered as sister to *Epistephium* (LPP = 0.89) (Figs. 2, 3b). Most remaining topological differences were confined to subgenus *Vanilla*, particularly in the placement of taxa comprising the Palmarum grade: in the Nuclear_ML analysis, *V. bicolor* was recovered as the earliest diverging lineage (BBS = 87), followed by a clade comprising *V. palmarum* and *V. sprucei* (BBS > 90), whereas in the Nuclear_Astral analysis *V. palmarum* was recovered as the first diverging lineage (LPP > 0.99), followed by *V. sprucei* (LPP = 0.62) and *V. bicolor* (LPP = 0.36). The positions of *V. insignis* and *Vanilla_spnov_114* also differed markedly between the two analyses, with the Nuclear_Astral tree placing *V. insignis* within clade J and *Vanilla_spnov_114* within clade I, whereas the Nuclear_ML analysis recovered these two taxa as closely related and sister to the clade comprising clades I and J.

**Fig. 3.**
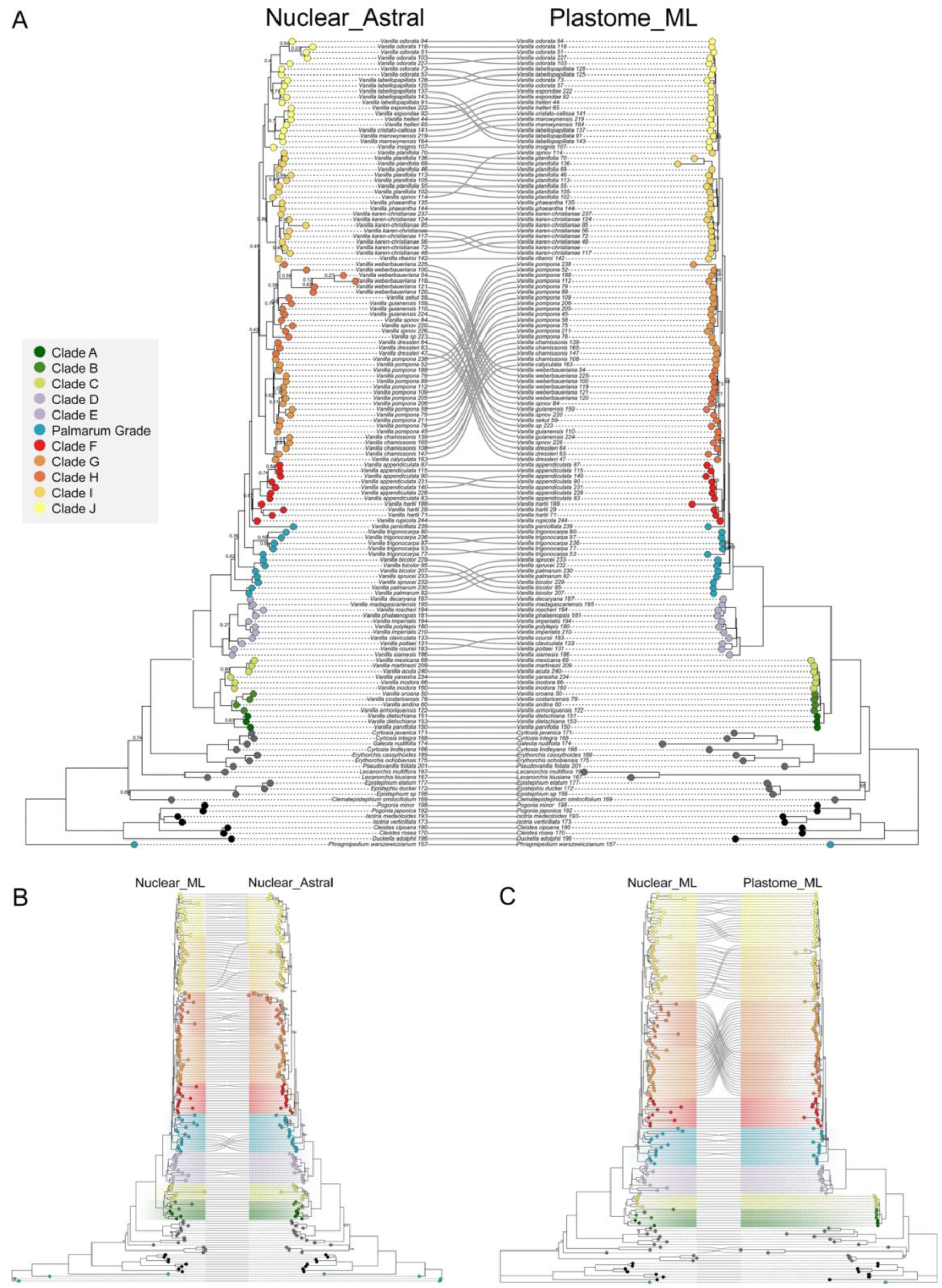
Phylogenetic tanglegram comparing (A) the nuclear ASTRAL species tree and the plastome maximum likelihood (ML) tree; (B) nuclear ML and ASTRAL trees; and (C) nuclear and plastome ML trees. Tip labels and colored boxes correspond to those in Fig. 2. Branch support values are shown only for LPP < 0.9 in the ASTRAL tree and BS < 90 in the plastome trees. Only taxa shared between the compared datasets were retained.

A comparison between the Nuclear_Astral and the Plastome_ML reveals broadly similar relationships but also noteworthy topological conflicts (Figs. 3a, S5). Within tribe Vanilleae, *Lecanorchis* is sister to the remainder of the tribe (LPP = 0.74) in the MSC tree (Figs. 3a, S1-2), whereas in the Plastome_ML tree it appears more closely related to a clade comprising *Epistephium* and *Clematepistephium*, although this relationship is poorly supported (BBS = 55) (Figs. 3a, S4). Both analyses strongly support the three major *Vanilla* clades (LPP = 1, BBS = 100). Differences are observed within *Vanilla* subg. *Membranacea*, where Clade B is monophyletic in the Nuclear_Astral tree but paraphyletic in the Plastome_ML tree. Relationships within *Vanilla* subg. *Thetyos* are consistent across both datasets. The greatest discordance occurs in *Vanilla* subg. *Vanilla*, beginning with the *Palmarum* grade. In the Nuclear_Astral tree, *V. palmarum* is sister to the remainder of the Palmarum grade followed by *V. sprucei* and *V. bicolor*, whereas in the Plastome_ML tree the order is reversed (*V. bicolor*, then *V. palmarum*, then *V. sprucei*). Relationships among the recognized Clades F, G and H also differ between analyses. In the Nuclear_Astral tree, these form a grade that is sister to a clade comprising clades I and J (LPP = 0.43–0.49), whereas in the Plastome_ML tree, clade F is sister to clade H (BBS = 100), and together they are sister to the remaining members of subg. *Vanilla*, with clade G recovered as sister to the clade of clades I and J (BBS = 100). Consistent with this pattern, comparisons between Nuclear_ML and Plastome_ML topologies likewise indicate that the major discordances are concentrated in the relationships among clades G and H and in the placement of *Vanilla_spnov_114* and *V. insignis* (Fig. 3c).

Focal common nodes of discordance identified above are consistent with overall high genealogical conflict across multiple support metrics, including gene and site concordance factors (gCF and sCF), normalized quartet support (QS), and the polytomy test (Figs. S6-9). Our analysis revealed moderate overall gene-tree incongruence (normalized quartet score = 0.7591), with most genus-level backbone nodes exhibiting high concordance (gCF and sCF > 50, LPP = 1, QS ≈ 0.8, *p* = 0). The primary deviation involves the placement of *Lecanorchis* as sister to the remainder of *Vanilleae*, which displays low concordance factors (gCF = 18, sCF = 36.3), moderate LPP (0.74), and QS (0.4), although the polytomy test strongly rejects unresolved relationships (*p* = 0). *Vanilla* and its three major subgenera are highly supported across all metrics (gCF = 60–77, sCF = 42–78, LPP = 1, QS = 0.8–0.9, *p* = 0). However, substantial incongruence occurs at several inner nodes along the backbone, with the nodes giving rise to Clades G and H exhibiting very low concordance factors (gCF = 0–0.6, sCF = 34–42), low LPP and QS values (LPP = 0.43–0.49, QS = 0.34–0.35), and polytomy tests that fail to reject unresolved relationships (*p* = 0.4–0.95). Within subg. *Vanilla*, discordance is concentrated in the *Palmarum* grade, where *V. bicolor* and *V. sprucei* show the lowest concordance factors (gCF = 7–9, sCF = 27–33), moderate to low LPP (0.3–0.6), low QS (∼0.3), and polytomy test results consistent with unresolved branches (*p* ≈ 0.5–0.9). No section within subg. *Vanilla* are strongly supported by concordance factors (gCF = 0.8–18; sCF = 44–57), with clade H as an outlier (sCF = 74.2), contrasting with generally high LPP values (0.9–1) and moderate QS (∼0.4, reaching 0.66 in clade H). Most crown nodes reject the polytomy hypothesis (*p* = 0), except for clade F (*p* = 0.332).

The network analysis conducted using SplitsTree recovered the three major clades of *Vanilla* as well as the five subclades within subgen. *Vanilla* (Figs. 1A, S10). However, considerable topological conflict was detected among focal clades, as evidenced by the short internal branches and strong interconnections among several minor groups. These conflicts are particularly pronounced among the three clades of *Vanilla* subgen. *Membranacea*, the two clades of *Vanilla* subgen. *Thethyos*, and within the Palmarum grade of *Vanilla* subgen. *Vanilla*.

Our analysis testing the null hypothesis of equal proportions of the two discordant quartets for each node based on nuclear gene trees revealed that several nodes are well-resolved, as indicated by low ILS and IH indices, including all nodes within Pogonieae (ILS-i ≈ 0.3%, IH-i ≈ 0.3%), the crown node of Vanilleae (ILS-i = 6%, IH-i = 0%), and the clade of fully mycoheterotrophic vanilloids (e.g., *Pseudovanilla*, *Galeola*) (ILS-i = 9%, IH-i = 0%). *Vanilla* and its subgeneric taxa were also recovered as highly resolved (ILS-i = 0.05–0.3%, IH-i = 0–0.3%) (Figs. 4, S11). Strong signals of ILS were detected within *Vanilla* subg. *Vanilla* (ILS-i > 0.7%, *p* > 0.1), likely explaining the low support values observed in concordance factors, QS, and the polytomy test at these nodes. Similar ILS signals were observed at inner nodes of *Vanilla* subg. *Membranacea*, particularly within Clades B and C (ILS-i = 0.74–0.9%), and within *Vanilla* subg. *Thethyos*, for example at the basal node leading to Antillean *Vanilla* and its African relatives (ILS-i = 1%). Conversely, significant signals of introgressive hybridization were detected at five nodes, including the stem nodes of *Lecanorchis* and the mycoheterotrophic vanilloids (IH-i = 40–47%, *p* < 0.01), the most recent common ancestor (MRCA) of Clades B and C (IH-i = 21%, *p* < 0.05), the clade comprising *V. atropogon* Schuit., Aver. and Rybková and *V. pilifera* Holttum within Clade D (IH-i = 23.5%, *p* < 0.05), and the node connecting *V. phaeantha* with a taxon endemic to the Yucatán (*Vanilla_spnov_114*) (IH-i = 32%, *p* < 0.01).

**Fig. 4.**
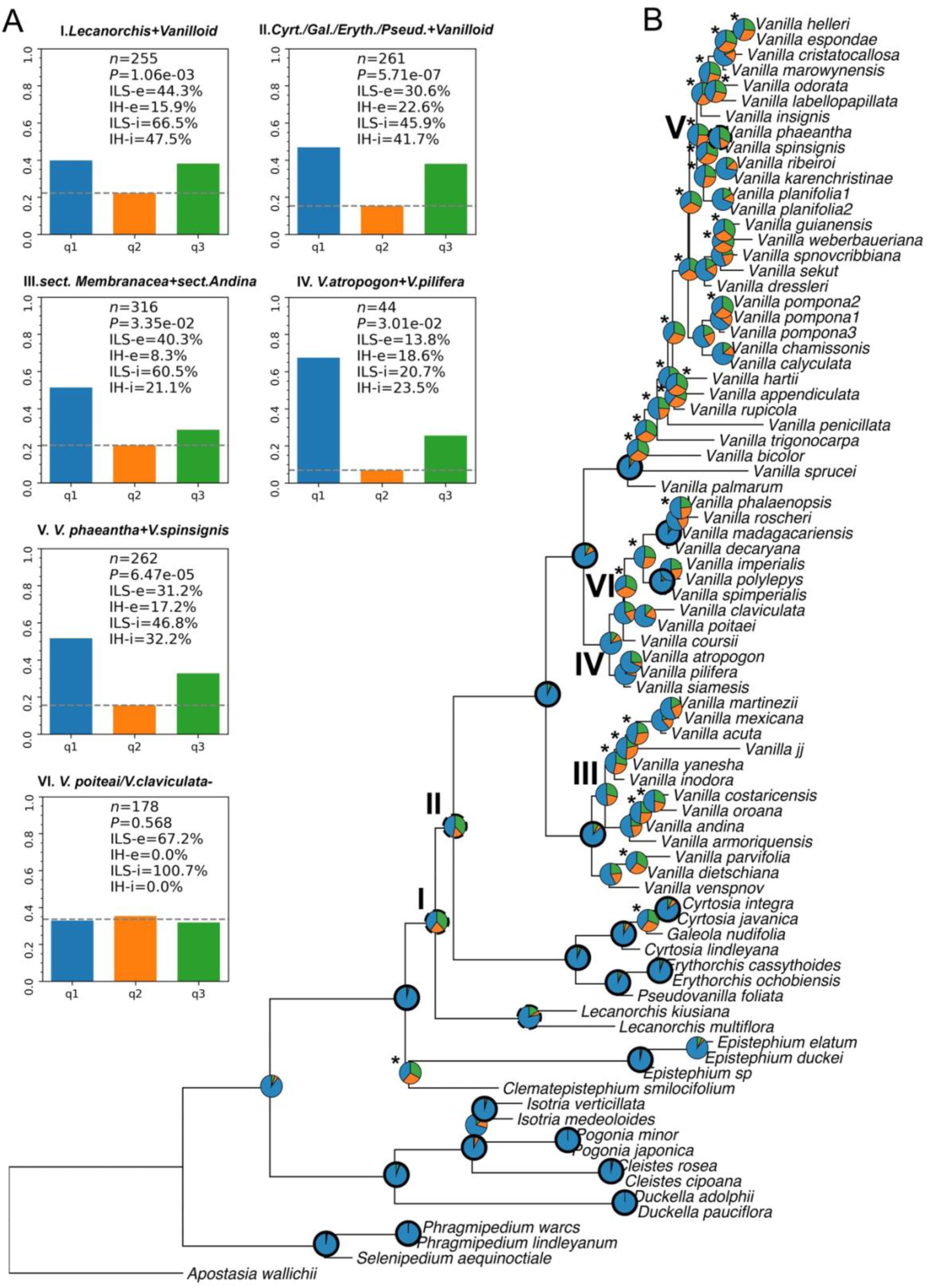
Detection and characterization of gene-tree discordance through visualization of topological conflicts between nuclear genes and the species tree, as quantified using *PhyTop*. (A) Tests of deviation from the expectations of ILS, where the null hypothesis is *q₂ = q₃*. Roman numerals indicate cases with IH-i > 20% (I–V), and VI marks an extreme case with ILS-i = 100%; (B) ASTRAL species tree including one representative tip per species; pie charts at each node show the proportion of concordant quartets (light blue) and the two discordant quartets (orange and green). Each box in (A) includes the following metrics: *n* = number of gene trees for that node; *P* = *p*-value for rejecting the null hypothesis; ILS-e = percentage of loci supporting ILS; IH-e = percentage of loci supporting introgressive hybridization (IH; alternative hypothesis); ILS-i = index of ILS; IH-i = index of IH. Asterisks in the ASTRAL tree (B) denote nodes with ILS-i > 70%, bold circles indicate well-resolved nodes (ILS-i ≈ 0.3%, IH-i ≈ 0.3%), and dashed circles indicate *p* < 0.03.

Partitioned phylogenetic network analyses at the genus level identified two reticulation events (−logL = −14362.49). In the optimal network, *Lecanorchis kiusiana* inherited 73.5% of its ancestry from *L. multiflora* and 26.5% from *Galeola nudifolia*. Moreover, the clade comprising all mycoheterotrophic vanilloids and *Vanilla* likely originated via ancient hybridization involving the ancestor of Vanilleae (15.4%) and *Lecanorchis* (84.6%) (Fig. S12). Within *Vanilla*, we detected a total of five hybridization events. In subgen. *Membranacea*, two events were supported (-logL=32093.19): Clade B appears to have arisen through ancient hybridization between *Vanilla_spnov_212* (21.7%) and *V. inodora* Schiede (78.3%), and the clade of *V. martinezii*, *V. mexicana*, and *V. guianensis* likely originated from *V. inodora* (73.8%) and *Vanilla yanesha* Damian (26.2%) (Figs. 5a. S12). Moreover, in the subg. *Thethyos* a single event was supported (-logL= 37034.21) with the clade made of *V. imperialis* Kraenzl. and *V. polylepis* Summerh. most likely originated by the MRCA of Clade E (excluding the caribbean taxa) (29.3%) and a similar looking *Vanilla imperialis (V_imperialis_184*) (Figs. 5b, S12). Within subgen. *Vanilla*, two hybridization events were inferred, including the clade of *V. cristato-callosa*, *V. marowynensis*, *V. helleri*, and *V. espondae*, which likely originated from the MRCA of clades J and I (17.7%) and the MRCA of *V. odorata* and *V. labellopapillata* (82.3%) (Figs. 5c, S12). Additionally, a taxon endemic to the Yucatán (*Vanilla_spnov_114*) appears to have a hybrid origin between *V. insignis* (53.1%) and *V. phaeantha* (46.9%).

**Fig. 5.**
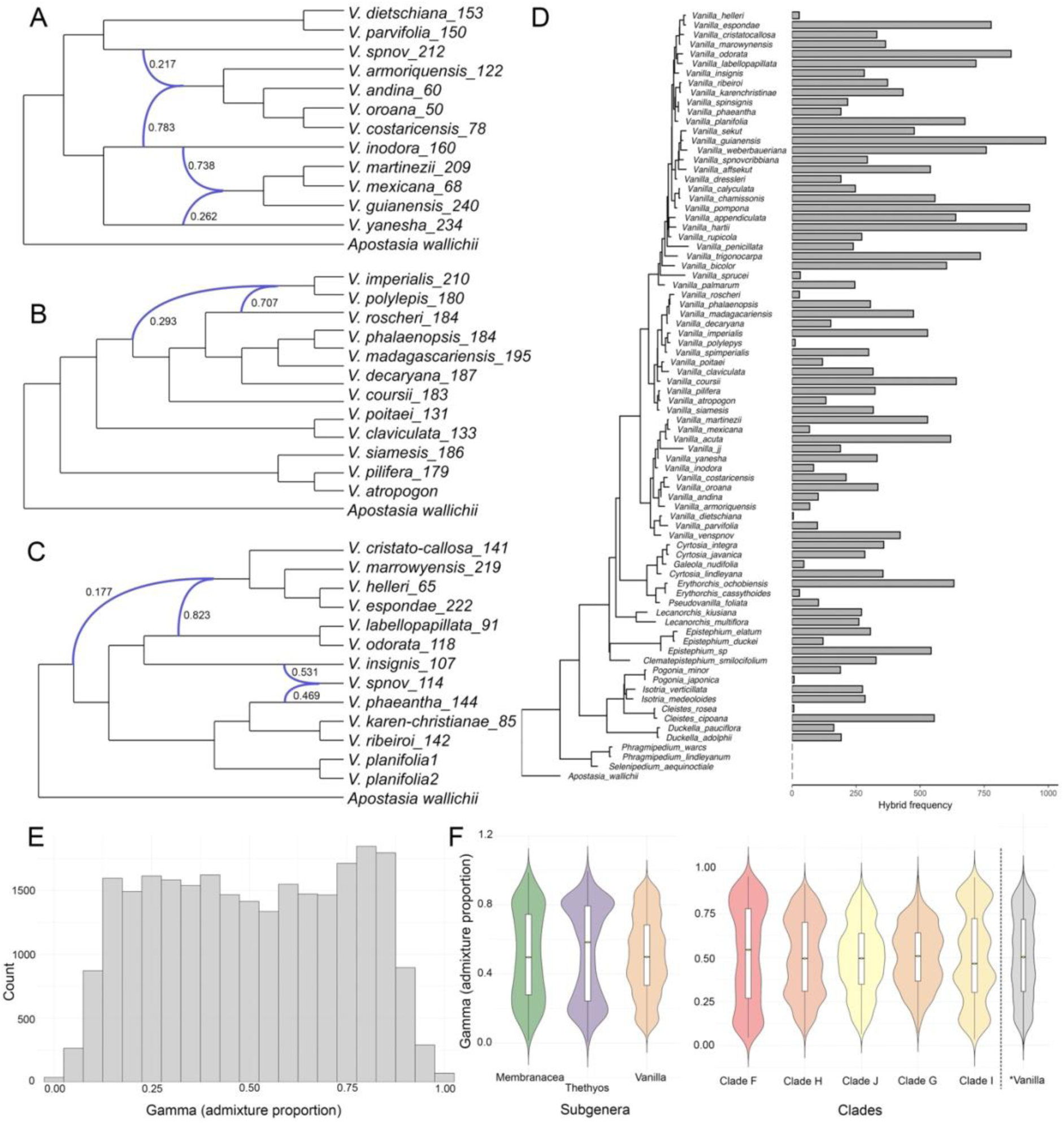
Interspecific gene flow detected using *PhyloNet* and *HyDe* analyses.(A–C) Phylogenetic networks reconstructed with *PhyloNet*, with curved blue arrows indicating hypothesized directions of ancestral gene flow and numerical values representing estimated proportions of genetic contribution between lineages. (D–F) Results of *HyDe* analyses: (D) ASTRAL tree shown alongside a bar plot summarizing the frequency with which each taxon is inferred as a hybrid in individual hybridization tests; (E) Overall histogram of γ values for the complete dataset; (F) Violin plots showing the distribution of γ across \**Vanilla* and within subgenera and major clades.

Analyses using *HyDe* partially corroborated *PhyloNet* findings, revealing 25931 triplets with significant signals of hybridization across the dataset (*p* < 0.05, *z-score* > 3), with most hybrid taxa detected within *Vanilla* subg. *Vanilla* and reaching the highest number of significant events in *V. guianensis* (Figs. 5d, S13-14). The distribution of γ values across the entire dataset was bimodal, suggesting an overall presence of ancient hybridization signals (Fig. 5e). At the tribal level, *Pogonieae* also exhibited a bimodal γ distribution, whereas most *Vanilleae* hybrids clustered around γ = 0.8 (Fig. S15). Overall, *Vanilla* displayed heterogeneous γ values without a consistent trend (Fig. 5f). Within the three major *Vanilla* subgenera, *Vanilla* subg. *Vanilla* showed a distribution similar to that observed at the genus level, whereas *Vanilla* subg. *Membranacea* and subg. *Thethyos* exhibited distributions skewed toward the extremes (γ ≈ 0 or 1), consistent with patterns of ancient hybridization (Figs. 5f). At the inner level, Clades F and H showed signatures of ancient hybridization (γ biased toward 0 and 1), while Clades J and G had γ values centered near 0.5, suggesting more recent hybridization events (Fig. 5f). Clade I exhibited a complex pattern, with up to four distinct peaks in its γ distribution. Finally, using *VisualHyDe*, we identified six significant hybridization events at the clade level: (a) involving the MRCA of *V. labellopapillata* and *V. odorata* (γ ≈ 0.5); (b) *V. phaeantha* and *Vanilla_cf_114* (γ ≈ 0.2–0.8); (c) *V. guianensis* and *V. sekut* (0 ≤ γ ≤ 1); and (d) *V. hartii* and *V. appendiculata* (0 ≤ γ ≤ 1). The remaining two events occurred in *Lecanorchis* and *Isotria*, both exhibiting 0 ≤ γ ≤ 1 (Fig. S16).

### Divergence time and biogeography

Our analyses revealed that the best-fitting model varied between DEC and DIVALIKE across different settings, with substantial differences between the time-stratified and unconstrained configurations. Because the time-stratified approach more accurately reflects the paleogeographic history of the Americas, all subsequent interpretations are based on this framework (Table S4). For the complete dataset, which included all vanilloid lineages, the best-supported model was DIVALIKE+J (lnL = –61.87, AIC = 129.7). This model inferred a Neotropical origin for Vanilloideae during the Late Cretaceous, with an estimated crown age of 77.2 Ma (95% HPD: 75.48–78.89 Ma) (Figs. S17, S18). Both Pogonieae and Vanilleae originated and diversified in the Neotropics during the Paleocene, with crown ages of 59.89 Ma and 65.28 Ma, respectively (95% HPD: 57.99–61.79 Ma and 63.62–66.9 Ma). The MRCA of Vanilleae dispersed into the Nearctic during the Eocene (43.8 Ma; 95% HPD: 42–46 Ma), later expanding into the Palearctic in the Oligocene (27.2 Mya; 95% HPD: 16–38 Ma). This expansion was followed by a vicariant split that resulted in *Cleistesiopsis* Pansarin and F.Barros and *Isotria* Raf. being restricted to North America and *Pogonia* to Eurasia. Among the non-*Vanilla* Vanilleae, two early Paleocene long-distance dispersal events were inferred from a Neotropical MRCA: (a) to the Palearctic and Indo-Malayan regions, giving rise to *Lecanorchis* Blume, and (b) to Australasia and Indo-Malaya, leading to the holo-mycoheterotrophic vanilloids, which began diversifying in the late Oligocene (24.7 Ma; 95% HPD: 15–35 Ma). The MRCA of this holo-mycoheterotrophic group subsequently underwent a vicariant split, separating a clade restricted to Indo-Malaya (*Cyrtosia* Blume and *Galeola* Lour.), including a later back-dispersal into Australia by *Cyrtosia javanica*, from a second clade composed of *Erythrorchis* Blume and *Pseudovanilla* Garay, both endemic to Australasia.

Within our Vanilla-only time-stratified analysis, DEC+J was the best-supported model (lnL = –228.6; AIC = 463.6), inferring a Guiana Shield origin for *Vanilla* and an onset of diversification in the Early Oligocene (29.8 Mya; 95% HPD: 18–40 Mya) (Figs. 6, S17). The earliest divergence separated *Vanilla* subg. *Membranacea* from a large clade comprising Vanilla subg. *Thethyos*, Vanilla subg. *Gondwana*, and *Vanilla* subg. *Vanilla*. Subg. *Membranacea* subsequently underwent a short-range dispersal into Amazonia, whereas the MRCA of the larger clade retained a French Guianan ancestral range. The ancestor of this large lineage began diversifying in the early Miocene (20.63 Ma; 95% HPD: 12.7–28.7 Ma), later giving rise to *Vanilla* subg. *Vanilla*, which likely had a broadly South American–Antillean distribution despite uncertainty in ancestral state reconstruction. Its sister lineage (*Vanilla* subg. *Thethyos* + *Vanilla* subg. *Gondwana*) retained the Guiana Shield range. Within this lineage, three long-distance dispersal (LDD) events were inferred: (i) from the Guiana Shield to Africa in the mid-Miocene, giving rise to the ancestor of Old World taxa; (ii) from Africa to Eurasia in the late Miocene, forming the exclusive Asian lineage (Clade D); and (iii) from the MRCA of the African lineage (Clade E) to the Antilles during the late Miocene (8.9 Mya; 95% HPD: 5.47–13 Mya).

**Fig. 6.**
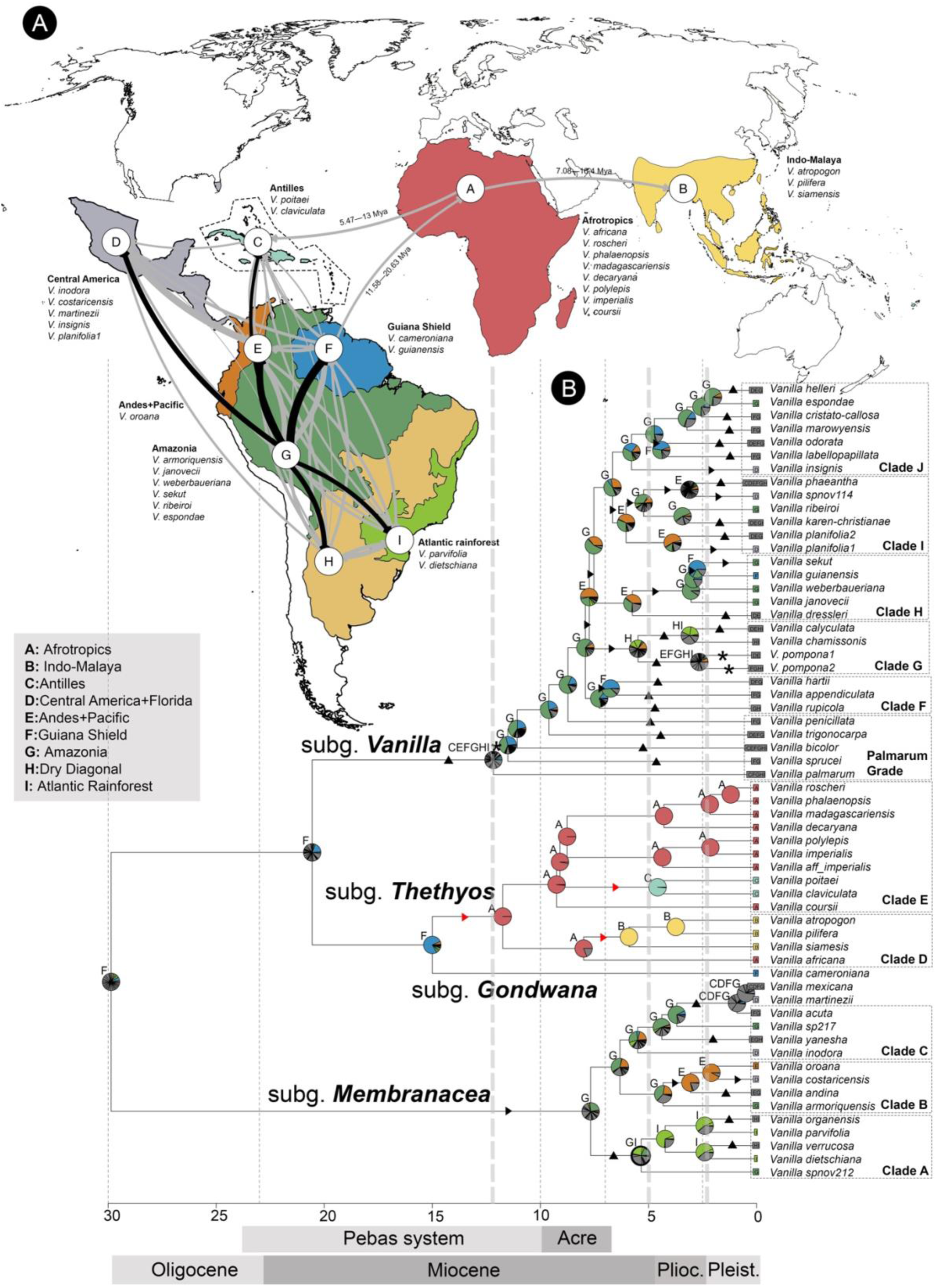
Historical biogeography of Vanilla. (A) Dispersal patterns modeled from 50 biogeographical stochastic maps of Vanilla, displayed on a world map using the DEC+J model. Arrows indicate the direction and frequency of dispersal events: black arrows highlight dispersals from Amazonia, while grey arrows represent all other movements. Arrow thickness reflects the total number of events. Divergence-time estimates for major long-distance dispersal events are indicated. The map is not to scale. (B) Dated phylogeny of *Vanilla* with ancestral ranges inferred using maximum likelihood under the best-supported DEC+J time-stratified model. The timescale is in millions of years before present. Pie charts at each node depict the relative likelihood of alternative ancestral ranges, with letters indicating the range with the highest probability. Thickened circles mark inferred vicariant events; red triangles indicate long-distance dispersal events; black triangles denote range expansions; arrowheads represent dispersal events; and asterisks indicate reductions in ancestral range size. Grey arrows indicate periods of accelerated Andean uplift (Gregory-Wodzicki, 2000).

Among exclusively Neotropical lineages, Vanilla subg. *Membranacea* began diversifying in the late Miocene (7.82 Mya; 95% HPD: 4.7–10.7 Mya). The MRCA of Clade A dispersed from Amazonia to the Atlantic Forest, followed by range expansions into the Dry Diagonal in *V. organensis* and *V. verrucosa* at the Pliocene–Pleistocene boundary, and a back-dispersal to Amazonia in *Vanilla* sp. nov. 212. The MRCA of Clades B and C retained an Amazonian range. Clade B dispersed to the Northern Andes + Pacific during the Pliocene (4.3 Mya; 95% HPD: 2.19–6.34 Mya), later reaching Central America (*V. costaricensis*) and expanding back into Amazonia (*V. andina*) during the Pleistocene. Clade C’s ancestor remained Amazonian, dispersed to Central America in the late Miocene (*V. inodora*), expanded into the Northern Andes and Dry Diagonal in *V. yanesha*, and underwent a major expansion to the Antilles, Central America, and the Guiana Shield in the MRCA of *V. acuta*, *V. mexicana*, and *V. martinezii* during the Pliocene (3.63 Mya; 95% HPD: 2.32–5 Mya).

In the vanillin-producing clade (*Vanilla* subg. *Vanilla*), most diversification occurred within the last 10 Myr, with a pronounced burst during the Late Miocene (∼7 Ma). The early ancestor occupied a broadly inferred South American range, which later contracted to the Guiana Shield, with a short mid-Miocene dispersal into the Amazonia (11.92 Mya; 95% HPD: 7.55–15.03 Mya). The deepest-diverging taxa of the *Palmarum* grade underwent multiple range expansions from the Guiana Shield and Amazonian early nodes, while *V. palmarum* retained much of the ancestral range and expanded into the Antilles. Similarly, *V. bicolor* dispersed widely from an Amazonian ancestor across South America and the Antilles. *V. trigonocarpa* expanded from the Amazon into Central America, the Northern Andes + Pacific, and the Guiana Shield, whereas *V. penicillata* and *V. sprucei* dispersed only from the Amazon to the Guiana Shield.

Within *Vanilla* subg. *Vanilla*, major subclades experienced multiple dispersal and range expansion events, mostly during the late Miocene and Pliocene. Dispersal primarily occurred across internal nodes, involving the Amazonian region and the Northern Andes + Pacific, with fewer events involving the Guiana Shield. Most MRCAs of major lineages lost their Amazonian range, except for Clades F and J, while Clades H and I were inferred to have a Northern Andes + Pacific ancestor, and Clade G was confined to the Dry Diagonal. Notable range expansions include the MRCA of *V. phaeantha*, *V. ribeiroi*, and *V. karen-christianae*, which spread from the Amazon to all South American regions defined in this study, and the MRCA of Clade G, which expanded from the Dry Diagonal into the Atlantic rainforest, giving rise to the *V. calyculata* and *V. chamissonis* clade. Although many expansions occurred in the late Miocene and Pliocene, several are inferred to have taken place more recently, within the last ∼5 Mya at the Pliocene–Pleistocene boundary. Some involved minor range increases, such as *V. dressleri* (Northern Andes → Central America) and *V. appendiculata* (Guiana Shield → Amazon), whereas others, notably *V. pompona* and *V. phaeantha*, expanded extensively, now occupying six and seven of the nine Neotropical regions defined in this study, respectively.

Our biogeographical stochastic mappings (BSMs) revealed that most events were within-area speciation (32%) or dispersal (64%), with vicariance being rare (4%) (Table S9). Vicariant events occurred primarily within an ancestral range comprising the Northern Andes + Pacific and Amazonia, with the highest mean number of events not exceeding 0.78 (Table S12). Among dispersal events, range expansions predominated (mean = 71) over founder events (mean = 12), and dispersal patterns varied considerably among regions (Table S10). Amazonia was the primary source (31%), followed by the Andes + Pacific (17%), mostly dispersing to the Guiana Shield and Central America. The most frequent movements were Amazonia → Guiana Shield (11 of 83 events) and Andes + Pacific → Central America (8 of 83), while other transitions, such as Amazonia → Central America or Amazonia → Andes + Pacific, occurred with moderate frequency (Table S10). Dispersals among New World regions accounted for 98% of events, whereas Old World or transcontinental dispersals were infrequent (2.4%), with the Afrotropics serving as a source for Indo-Malaya and the Antilles. Range expansions largely mirrored overall dispersal patterns, emphasizing Amazonia as the dominant source and Central America as the primary sink (Fig. S20, Tables S10, S11), whereas founder events were dominated by Amazonia → Andes + Pacific transitions, with other founder-event dispersals occurring more evenly among regions. Overall, dispersal movements were highly symmetric (∼86% of area pairs), such as between the Dry Diagonal and Atlantic Rainforest (2.92 ± 1.83 vs. 3.76 ± 1.53), but notable asymmetries existed, most prominently Andes + Pacific → Central America, which occurred ∼38 times more frequently than in the reverse direction (Table S10).

## DISCUSSION

### Novel insights into the evolution of Vanilla and its allies

Our analysis recovered all major infrageneric *Vanilla* taxa recently proposed by our research group as monophyletic (Karremans *et al.*, 2025), including the monotypic *Vanilla* subgen. *Gondwana*. Notably, this subgenus is placed phylogenetically for the first time as sister to the Old World *Vanilla* subgen. *Thethyos* based on our plastome dataset (Fig. S4). At the sectional level, we sampled six of the seven sections recognized in this framework (excluding the African *sect. Thethyos*) and recovered four as monophyletic, including *Vanilla sect. Dictyophyllaria* and *Vanilla* sect. *Membranacea* within *Vanilla subg. Membranacea*, and *Vanilla sect. Aphyllae* and *Vanilla* sect. *Miguelia* within *Vanilla subg. Thethyos*. Although sampling of the Old-World clade (*Vanilla* subg. *Thethyos*) remains limited, our analyses place the Antillean taxa, nested within this predominantly Old-World subgenus, in a deeper position, sister to all of *Vanilla* sect. *Aphyllae*. This topology contrasts with previous reconstructions, which positioned them shallowly within the section (Bouétard, 2010; Givnish *et al.*, 2016).

The two recently proposed sections for vanillin-producing clade (*Vanilla subg. Vanilla*: *Vanilla sect. Palmarum* Karremans, Damián & Pupulin and *Vanilla sect. Vanilla*) were not recovered as monophyletic in our genome-based analysis. Instead, we identified a paraphyletic assemblage at the deepest node and five highly supported subclades. The basal grade, termed the “Palmarum grade”, partially overlaps with *Vanilla sect. Palmarum* and includes *V. palmarum*, *V. sprucei*, *V. bicolor*, *V. trigonocarpa*, and *V. penicillata*. Notably, the position of *V. penicillata* within this grade was unexpected, as it had previously been considered morphologically allied to *V. ribeiroi* (Soto Arenas & Cribb, 2010). This basal grade is highly heterogeneous and exhibits unusual morphological traits relative to the remainder of the subgenus, including epiphytic habit, non-vanillin producing fruits (*V. palmarum*, *V. bicolor*); scale-like or non-coriaceous leaves (*V. penicillata*, *V. trigonocarpa*, *V. sprucei*), and branching inflorescences (*V. sprucei*, *V. palmarum*).

Within the remainder of *Vanilla subg. Vanilla*, we identified five well-supported subclades, which we identified as Clades A-J. Affinities within some of these groups had been suggested previously, particularly by Soto Arenas and Cribb (2010), who defined six “informal groups” under their *Vanilla sect. Xanata* (now *Vanilla subg. Vanilla*) based on morphological similarities. Our analysis recovered two of these groups as monophyletic (clades G and H), whereas the remaining taxa were distributed among the other four subclades or included in the Palmarum grade. Although several attempts have been made over the last two decades to resolve relationships within *Vanilla subg. Vanilla*, the presence of poorly supported clades and polytomies has hindered a clear understanding of its evolutionary history (e.g., Pansarin & Fernandez 2023; Pansarin, 2024; Karremans *et al.*, 2025). Our results provide a more robust phylogenetic resolution, with most internal nodes strongly supported and clarify the positions of many previously unplaced taxa, including the close relationships among *V. espondae*, *V. cristato-callosa*, and *V. marrowynensis*, as well as *V. karen-christianae* and *V. ribeiroi*. Nevertheless, some relationships remain unresolved and weakly supported, particularly within Clade H, and along several backbone nodes leading to Clades G and H, which are characterized by short branches and low phylogenetic resolution.

With respect to the membranaceous *Vanilla* (*Vanilla* subg. *Membranacea*), our analysis is largely congruent with previous inferences based on nrITS (Pansarin, 2024, 2025; Krahl *et al.*, 2025) and with our recent supermatrix analysis incorporating *nrITS, xdh, rbcL, matK, psaB, psbB, psbC, trnL, atpA*, and *nad1* (Karremans *et al.*, 2025). In particular, Clade B is more extensively sampled here, now including at least four species compared with only two (*V. sarapiquensis* and *V. costaricensis*) in previous work. However, the placement of *V. sarapiquensis* within Clade B requires further verification, as it was not sampled in this study and is morphologically more similar to members of Clade C, especially *V. mexicana*. While most internal relationships within *Vanilla* subg. *Membranacea* were well supported in previous studies (e.g., Pansarin & Fernandes, 2023; Pansarin, 2024), the relationships among the major clades, especially Clades C and A, the only widely sampled representatives, have historically been weakly supported and contentious. Our results help clarify these uncertainties, showing that, although Clades B and C are closely related, they do not form a single clade. Instead, Clade B is sister to Clade C, and together they are sister to Clade A, with all nodes strongly supported. In addition, we identify a likely undescribed taxon, morphologically similar to members of Clade A, occurring in the Venezuelan Caribbean, a placement that is unexpected and further explored in the biogeographical analysis.

### Phylogenetic placement of the closest wild relatives of Vanilla planifolia

A major outcome of our phylogenomic analyses is the resolved placement of *V. planifolia*, the primary species used for natural vanillin production, and the precise identification of its closest wild relatives (CWR). Previous studies have struggled to resolve the position of *V. planifolia* within the *Vanilla* phylogeny, frequently recovering it in poorly supported or unresolved topologies. Early analyses based on *nrITS* and *rbcL* data variously placed *V. planifolia* within a clade including *V. insignis* and *V. pompona* (Soto Arenas, 2003), as sister to *V. phaeantha* (Cameron, 2010; Bouétard et al. 2010), or in poorly supported polytomies (Soto Arenas & Dressler 2010; Azofeifa et al. 2017; Pansarin 2024, 2025). More recently, genomic studies, leveraging the draft genome of *V. planifolia* and thousands of SNPs generated through genotyping-by-sequencing have likewise suggested a close relationship with *V. phaeantha* (Hu *et al.*, 2018; Chambers *et al.*, 2021).

Our genome-scale analyses, combined with expanded taxon sampling, place *V. planifolia* within Clade I that includes *V. karen-christianae*, *V. ribeiroi*, *V. phaeantha*, and a taxon endemic to the Yucatán (Fig. 1b), thereby confirming and refining earlier hypotheses of affinity with *V. phaeantha*. This phylogenetic resolution has important applied implications. Commercial vanilla cultivation relies predominantly on vegetatively propagated descendants (i.e., clones) of *V. planifolia*. As a result, the cultivated gene pool exhibits extremely limited genetic diversity (Bory *et al.*, 2008; Minoo *et al.*, 2008). This genetic uniformity constitutes a substantial risk to global supply chains, as clonally propagated crops tend to be uniformly susceptible to major pathogens, including *Fusarium* species and several viral diseases (Koyyappurath *et al.*, 2016). In this context, the robust identification of the closest wild relatives of *V. planifolia* is critical for broadening the genetic base of cultivated vanilla, mitigating production risks, and strengthening the long-term resilience and sustainability of global vanilla production systems (Watteyn *et al.*, 2025; Serna-Sánchez *et al.*, 2025).

Moreover, our analyses of eight *V. planifolia* accessions reveal a clear population structure comprising two strongly supported subclades: one endemic to northern Mesoamerica (Mexico, Guatemala, and El Salvador) and another extending from Central and South America (Costa Rica to Peru). These distributions correspond closely to established biogeographical divisions, with northern populations ocurring within the Mesoamerican dominion and southern populations within the Pacific and South Brazilian dominions (sensu Morrone, 2014). Although clear genetic and biogeographic differentiation exists between northern and Central-Southern populations, morphological differences are minor. A thorough phenotypic assessment is needed to determine whether these differences are diagnostic enough to support separate species or if a subspecies classification is more appropriate.

### Phylogenetic incongruence in Vanilla evolution

Based on our analyses, we reconstruct a robust evolutionary framework for *Vanilla*, with nodes at the genus, subgenus, and species levels generally well supported, showing LPP values close to or equal to one. However, CFs revealed notable conflict among several gene tree topologies. Although the three major *Vanilla* subgenera are highly congruent, the shallowest inner nodes exhibited substantially lower CFs, particularly within *Vanilla* subg. *Vanilla*, a pattern largely driven by extremely low gCF values. Deeper nodes leading to members of the *Palmarum* grade and within the backbone of *Vanilla* subg. *Vanilla* also showed similarly low CFs, consistent with lower QS values and with the polytomy test failing to reject resolved branches. Low quartet support and CFs paired with high LPP values, precisely the pattern observed here, have been reported in other orchid lineages (Zhang *et al.*, 2023; O’Donnell *et al.*, 2024) and specifically in datasets generated from the Angiosperms353 target-capture loci (Murillo-A. *et al.*, 2022; Barret *et al.*, 2025,). This inverse relationship is expected when heterogeneous shallow and deep nodes are present or when sampling spans broad taxonomic scales, leading to high LPP values that nevertheless reflect support from only a small subset of gene trees. Together with our *PhyTop* analysis, these results revealed a high proportion of variation attributable to ILS (Fig. 4; Fig. S11). Such scenarios commonly arise when internal branches are short, a hallmark of rapid radiations, as is evident in our dataset, particularly within the vanillin-producing clade (*Vanilla* subg. *Vanilla*), and are consistent with previous studies showing that ILS can hinder species-level phylogenetic resolution (Riggins & Seigler, 2012; Gagnon *et al.*, 2021), especially in rapid radiations (Stull *et al.*, 2023).

Our study constitutes the first explicit evaluation of cyto-nuclear incongruence in *Vanilla*, filling a long-standing gap that primarily reflects the scarcity of plastid-based phylogenetic hypotheses for the genus. Over the past decades, most *Vanilla* phylogenetic studies have relied almost exclusively on the nuclear nrITS (e.g., Pansarin and Menezes, 2023; Pansarin, 2024, 2025; Krahl *et al.*, 2025). In this context, our plastome data are congruent with the results of Bouétard et al. (2010), which were based on psaB, psbB, psbC, and rbcL, in recovering *V. leprieurii (= V. hartii)* and “*V. ensifolia*” CR0174 (likely not *V. ensifolia* but instead a species closely related to *V. cribbiana*), members of Clades F and H respectively, as forming a clade that is sister to a clade composed of Clade G and Clades I–J. This same relationship was also recovered by Cascales *et al*. (2023) using rbcL and matK, albeit with more limited sampling than in Bouétard *et al*. (2010). Overall, our nuclear and plastome phylogenies are largely congruent, with the main difference involving the relationships between Clades G and H. In addition, several taxa and internal relationships that are strongly supported in the Astral tree are recovered as non-monophyletic in the plastome analysis, including *V. odorata* and *V. weberbaueriana*, among others. This discrepancy in the plastome dataset likely reflects its lower levels of genetic divergence and its greater susceptibility to distortion through chloroplast capture coupled with hybridization and introgression (e.g., Li *et al.*, 2026; Pérez-Escobar *et al.*, 2021), which argues for placing greater weight on the nuclear Astral phylogeny when assessing evolutionary relationships in *Vanilla*.

Both our nuclear ML and Astral phylogenies recover largely the same topology and differ from the most recent phylogeny proposed by Pansarin (2025), which was based on nrITS, in three main respects: (a) the position of Clade F, which in our reconstruction is sister to the remainder of *Vanilla* subg. Vanilla (excluding the Palmarum grade), whereas in Pansarin (2025) it is sister to the clade comprising Clades H–I–J; (b) the failure to recover several species as monophyletic, including *V. planifolia, V. cribbiana, V. calyculata, V. odorata,* and *V. labellopapillata;* and (c) the lack of resolution of relationships within *Vanilla* subg. Thethyos. All of these results are inconsistent with the nuclear phylogenies presented here, which are mutually congruent, and suggest that the Angiosperms353 probe set yields reliable nuclear data at the species level for reconstructing evolutionary relationships across Vanilla.

Disagreement between nuclear- and plastid-based topologies could also be explained by ancestral gene flow (hybridization). However, in scenarios characterized by short internal branches and high levels of ILS, such events can be particularly difficult to detect. This is because the probability of gene flow among recently diverged lineages is elevated under these conditions (Hibbins and Hahn, 2021; Barret *et al.*, 2025b). Network-based approaches that explicitly test for ancient gene flow can nevertheless reveal the prevalence of historical reticulation, even in exceptionally rapid plant radiations. These methods provide a powerful framework for distinguishing introgression from stochastic ILS (Stull *et al.*, 2020; GPWG, 2024; Lagou *et al.*, 2024; Lin *et al.*, 2026). In our study, the first attempt to detect signals of introgressive hybridization using *PhyTop* revealed five significant rejections of non-ILS patterns, three of them within *Vanilla*, with two directly associated with cyto-nuclear incongruent nodes. Subsequent analyses using *PhyloNet* showed that these three nodes are involved in both ancient and more recent episodes of gene flow, with most events occurring over the past ∼10 Ma. Natural hybridization has long been recognized as an important driver of plant diversification and speciation (Soltis & Soltis, 2009). This appears to be the case in *Vanilla* as well, although to a lesser extent than the pervasive influence of ILS. While numerous *Vanilla* hybrids have been artificially produced, primarily for crop improvement and breeding programs (Chambers, 2018; Menchaca, 2018; Grisoni & Nany, 2021), only two cases of natural hybridization have been formally documented: one in the Antillean species *V. claviculata* and *V. barbellata* Rchb. f. (Nielsen & Siegismund, 1999; Nielsen, 2000), and a more recent report between *V. phaeantha* and *V. pompona* (Pansarin, 2025).

Here, we report a third naturally occurring hybrid arising from crosses between *V. phaeantha* and *V. insignis*. This nothospecies has been known for at least 15 years, with the first attempt to clarify its identity made by Villanueva-Viramontes *et al*. (2017), who suggested it might represent an undescribed taxon based on ISSR and ITS data. More recently, Favre *et al*. (2022) reached a similar conclusion, noting a genetic affinity between this taxon and the known hybrid involving *V. bahiana* Hoehne (= *V. phaeantha*) and *V. insignis*. Our genome-scale dataset confirms these earlier suspicions, and we argue that this entity indeed represents a natural hybrid. Favre and colleagues additionally proposed a hybrid origin for two other Neotropical taxa using a Bayesian clustering approach. Accession CR03612, which clearly belongs to *V. karen-christianae*, was inferred to contain genetic contributions from *V. planifolia*, *V. pompona*, and *V. odorata*. Accession CR0116, representing *V. labellopapillata*, was suggested to result from hybridization between *V. pompona* and *V. odorata*. These results contrast with our findings, as none of our analyses, across three independent methods, recovered evidence for reticulation involving *V. karen-christianae* or *V. labellopapillata*. Even when exploring more complex reticulation scenarios (H = 3 or H = 4) in *PhyloNet*, which yielded log-probabilities comparable to the simpler model (H = 2), we detected no hybridization signals involving either taxon. This discrepancy likely stems from methodological differences, particularly the reliance on Bayesian clustering approaches such as STRUCTURE in Favre et al. These methods perform well for detecting admixture among very recently diverged populations (dozens of generations) but are known to perform poorly in cases of ancient hybridization (millions of years) and may even generate false positives under such conditions (Wang, 2017; Pang & Zhang, 2025), the most plausible explanation here. Additional support for this interpretation comes from the biological implausibility of natural hybridization between *V. odorata* and *V. pompona*, given their highly specialized pollination systems and the strong specificity of plant–pollinator interactions documented for these species (Serna-Sanchez *et al.*, 2025).

### Spatiotemporal evolution of Vanilla

Our ancestral-area reconstruction supports a Guianan origin of *Vanilla* in the Oligocene (ca. 30 Mya), offering new insight into the tempo and mode of diversification in the genus and representing the first study to explicitly evaluate biogeographic processes in *Vanilla* (Fig. 6). Few studies have addressed orchid biogeography at the family level using phylogenetic approaches and only a few have included *Vanilla*. Among these, our findings are broadly consistent with Bouetard *et al*. (2010) and Givnish *et al*. (2016), both of which inferred a Neotropical origin with HPD intervals overlapping ours. In contrast, a more recent study by Pérez-Escobar *et al*. (2024) recovered a Nearctic–Neotropical origin for the MRCA of *Vanilla*, although their divergence-time estimates (ca. 30 Mya) remain broadly congruent with our reconstruction. The differences between Givnish *et al*. (2016) and Pérez-Escobar *et al*. (2024) likely reflect variation in data sources and analytical frameworks. Givnish *et al*. (2016) relied on plastome data and Pérez-Escobar *et al*. (2024) used nuclear data, which can differ in evolutionary rates and may influence phylogenetic inference (Guo *et al.*, 2023). The studies also differed in their biogeographic approaches: Pérez-Escobar et al. (2024) implemented a DEC model incorporating several alternative topologies to account for phylogenetic uncertainty, whereas Givnish *et al*. (2016) compared multiple models, including time-stratified analyses informed by geological history. These analytical differences likely provide further explanation as to why our nuclear-based results diverge from Pérez-Escobar *et al*. (2024).

Sampling density and area coding can also influence ancestral reconstructions, even when using similar genomic datasets (Wang *et al.*, 2024). Our study presents the most densely sampled *Vanilla* phylogeny to date (57 species, versus 23 in Pérez-Escobar et al. 2024 and 13 in Givnish *et al.*, 2016). Additionally, although the overall biogeographic areas considered are broadly comparable across studies, differences in how the deepest clades particularly the long-branched *Vanilla* subg. *Membranacea* are coded may have a substantial impact on ancestral-area estimates. For example, *V. mexicana* was coded as occurring in both the Neotropics and North America by Givnish et al. and Pérez-Escobar *et al.*, whereas we coded it as exclusively Neotropical because its only remaining Nearctic presence (southern Florida) is floristically more similar to the Caribbean and Central America flora (Fritsch and McDowell, 2003,

Santiago-Valentin and Olmstead, 2004). In Givnish *et al.*, the inclusion of close allies of *V. mexicana* mitigated the impact of this coding choice and the MRCA of *Vanilla* was still inferred as Neotropical. However, Pérez-Escobar et al. sampled only a single representative of *Vanilla* subg. *Membranacea*, coded as occurring in both the Nearctic and Neotropics, which likely pulled their estimate of the *Vanilla* crown node toward a Nearctic–Neotropical origin.

Initial diversification of Guianan *Vanilla* appears to have occurred during a period of relative geological stability. By the Oligocene, when *Vanilla* first diverged, the Guiana Shield represented one of the most stable regions of northern South America, forming part of the Amazonian Craton—an ancient, tectonically stable continental core characterized by ancient bedrock and generally nutrient-poor soils (Quesada *et al.*, 2009). This contrasts with the geologically younger western Amazonia, strongly influenced by Andean tectonics and dominated by nutrient-rich, Andean-derived sediments (McClain and Nauman, 2008). Diversification within *Vanilla* accelerated during the Miocene, when major dispersal events occurred from the Guiana Shield to nearby and distant regions. One plausible explanation for this shift is the increasingly dynamic geological conditions affecting the Guiana Shield and Amazon Basin during this time, including uplift and exhumation events that exposed deeper rock layers through erosion—processes documented several times in the past ∼500 Myr, including a major phase during the Miocene (Japsen *et al.*, 2025). This interval coincides with the formation of the Amazonian lowlands and the modern Amazon Valley, where most contemporary *Vanilla* populations occur. In this context, the Guiana Shield may have initially functioned as a stable diversification core during the Oligocene and later, during the Miocene, as a source area facilitating biotic exchange with the emerging Amazonian lowlands. This early diversification, which generated much of the extant diversity during the first half of the Miocene, is consistent with biogeographical patterns observed in other Guianan taxa, where “taxon-pulse” diversification cores have been proposed for bats (Lim 2012) and plant groups such as Bromeliaceae (Givnish *et al.*, 2011) and Rapateaceae (Givnish *et al.*, 2004).

Although the Guiana Shield served as the source of several early *Vanilla* lineages, our analyses suggest that most recent diversification occurred primarily in Amazonia and, to a lesser extent, in the Northern Andes and the Guiana Shield. These regions form what has been termed the “triangle of diversity” (Vásquez-Restrepo *et al.*, 2024), which in our dataset also shows the highest number of biotic interchange events, consistent with patterns reported in other Neotropical groups (Antonelli *et al.*, 2018). While most defined biogeographic regions have acted as both sources and sinks, Amazonia is strongly supported as the principal source region for *Vanilla*, particularly during the last ∼10 Myr (late Miocene to present). During this interval, membranaceous *Vanilla* lineages (*Vanilla subg.* Membranacea) began diversifying as the Amazon Basin transitioned from the extensive lacustrine–wetland Pebas system to the predominantly fluvial Acre system (Fig. 6), generating substantial fragmentation between northern and southern Amazonia (Hoorn *et al.*, 2010; Albert *et al.*, 2018; Bicudo *et al.*, 2019). At the same time, the South American Dry Diagonal was forming, although permeable forested corridors connecting Amazonia and the Atlantic Forest likely persisted until its full establishment in the Pleistocene (Sobral-Souza *et al.*, 2015; Thode *et al.*, 2019). Our biogeographic reconstruction supports this scenario, particularly within Clade A, which experienced a late Miocene range expansion from Amazonia into the Atlantic Forest, followed by a vicariant event that produced a northern Amazonian lineage and a clade with an MRCA restricted to the Atlantic Forest, later colonizing the Dry Diagonal during the late Pliocene and Pleistocene. This pattern mirrors numerous studies documenting recurrent Amazon–Atlantic Forest connections underlying widespread biogeographic disjunctions (reviewed in Della and Prado, 2025). Diversification in the other two major membranaceous clades (Clades B and C) occurred after the Northern Andes reached elevations of ∼3000 m (Hoorn *et al.*, 2010), with most range expansions and dispersal events from Amazonia into the Northern Andes–Pacific lowlands occurring within the last ∼5 Myr.

Vanillin-producing taxa (*Vanilla* subg. *Vanilla*) exhibit diversification patterns broadly consistent with their membranaceous congeners, with most deep nodes restricted to Amazonia and most range expansions and dispersal events occurring within the last ∼10 Mya (Fig. 6). The main exceptions involve the deepest nodes of the group, corresponding to the Palmarum grade, particularly the crown node of the subgenus, whose ancestral range in our analyses suggests a major colonization event from the Guiana Shield into adjacent regions during the early stages of the Pebas system (Hoorn *et al.*, 2010). This during a time, where much of South America remained predominantly tropical and forested, including areas that are today represented by deciduous forests or tropical savannas in southern Brazil and northern Venezuela (Bradshaw *et al.*, 2025). Although the Pebas system has traditionally been interpreted as a barrier limiting dispersal and gene flow in terrestrial organisms (Wesselingh & Salo 2006; Antonelli *et al.*, 2009), more recent perspectives consider it a dynamic and permeable system that promoted diversification through repeated cycles of connectivity and isolation across Amazonia and the Andes (Hoorn *et al.*, 2022). This scenario appears consistent with the early diversification of subg. *Vanilla*, which is characterized by extensive colonization events across South America, as inferred for the MRCA of the entire subgenus and subsequent dispersals such as those leading to *V. bicolor* and *V. palmarum*, both of Amazonian origin but now widely distributed across South America and the Antilles.

A recurring pattern in our reconstruction of *Vanilla* biogeographic history is that the Andes did not constitute a strong barrier to biotic exchange. Instead, the permeability of the Amazon–Pacific corridor appears to have facilitated repeated dispersals and range expansions into Central America, highlighting the role of this region as a major sink in *Vanilla* diversification. For example, within subg. *Membranacea*, the Central American endemic *Vanilla inodora* is most plausibly explained by a dispersal from Amazonia to Central America that occurred during a period of major Andean uplift (Gregory-Wodzicki, 2000). Similarly, the present distribution of *V. costaricensis* in Central America is best explained by an initial dispersal from Amazonia into the Northern Andes–Pacific region in the MRCA of Clade B, followed by a subsequent dispersal into Central America, likely facilitated by the closure of the Isthmus of Panama—an event that promoted dispersal and diversification in numerous Neotropical plant lineages (Luebert & Weigend, 2014). Comparable dynamics, in which colonization of the Northern Andes preceded dispersal into adjacent regions, are also evident in the Central American endemic *V. martinezii* and in *V. mexicana*, a widespread species that also reached the Antilles and, in the case of *V. mexicana*, as far as Florida. Likewise, vanillin-producing species (subg. *Vanilla*) appear largely unaffected by Andean uplift, with multiple dispersals from Amazonia to distant regions—including the Antilles and Central America—occurring within the last ∼5 Mya (Fig. 6), potentially driven by climatic instability in eastern Amazonia during the Pleistocene (Cheng *et al.*, 2013; Wang *et al.*, 2017; Silva *et al.*, 2019).

The apparent permeability of the Andes in Vanilla history is likely linked to its unique seed dispersal mechanism. Direct migrations from Amazonia to the Pacific and Central America has been reported for other plant groups including bromeliads and ferns (Sanchez-Baracaldo, 2004; Givnish *et al.*, 2011), as well as other orchid lineages (Pérez-Escobar *et al.*, 2017a,b). While orchid dispersal is often attributed to the high potential of anemochorous seeds (Arditi & Abdul, 2000), Vanilla produces fleshy fruits with thick, crustose seeds (Cameron, 1996), suggesting that alternative dispersal syndromes must be involved. Seed-dispersal research in *Vanilla* remains limited and is primarily concentrated on a small subset of species (reviewed by Karremans *et al.*, 2023a). Among the most comprehensively documented examples, evidence from Central America indicates that certain taxa exhibit a multimodal dispersal syndrome in which seeds are transported via both endozoochorous and ectozoochorous mechanisms mediated by bees and rodents (Karremans *et al.*, 2023b). Comparable processes have recently been proposed for some South American species as well (Pansarin, 2024). Recent phylogenomic and biogeographic evidence shows that *Proechimys*, an important disperser of *V. planifolia* and *V. pompona*, originated in the Miocene, with its Amazonian–Andean clade diversifying around the Pliocene (ca. 4 Mya) and later dispersing between these regions near the Pliocene–Pleistocene boundary (ca. 2 Mya) facilitated by the lower and warmer conditions of the Huancabamba Depression (Antonelli *et al.*, 2009; Dalapicolla, 2019; Dalapicolla *et al.*, 2024). These findings suggest that major geographic barriers, including the Andes, may not have strictly limited the movement of *Vanilla* dispersers, providing a plausible explanation for the recurrent Amazon–Andean biogeographic connections observed in the genus following Andean uplift.

Although Amazonia clearly served as a major center of in-situ diversification for *Vanilla*, reconstructing the genus’s full biogeographical history remains challenging. The biome is a vast, heterogeneous mosaic shaped by tectonic and hydrological changes in its drainage basins, and many key palaeogeological timelines are still debated (Cracraft *et al.*, 2020). Yet despite these uncertainties, one consistent finding across studies is that Amazonian landscape dynamics have repeatedly acted as powerful drivers of allopatric speciation a pattern mirrored across numerous vertebrate (Ortiz *et al.*, 2021; Pereira *et al.*, 2021; Vásquez-Restrepo *et al.*, 2024;) and plant lineages (e.g. Carvalho & Lohmann, 2020; Magri *et al.*, 2025), and one that likely applies to *Vanilla* as well.

### Systematic and Biogeographic Implications for Vanilloideae

Our results provide new insights into the relationships among major vanilloid lineages and the Vanilloideae subfamily as a whole. At the genus level, we sampled all vanilloid genera except *Eriaxis*, and based on our nuclear dataset, most relationships are broadly congruent with previous studies (e.g., Cameron, 1996, 2003, 2009). All nodes within Pogonieae are recovered with high support across all metrics, and no evidence of ancient reticulation is observed. Our nuclear and plastid topologies are largely congruent and align with most previous studies, with the notable exception of *Cleistesiopsis*. Although we were unable to retrieve high-quality Angiosperms353 loci from *C. divaricata*, we recovered 70 plastid loci using the CAPTUS pipeline for our reconstruction. Based on these data, *Cleistesiopsis* is inferred as more closely related to *Isotria* than to *Pogonia*, contrasting with most studies that placed it nearer *Pogonia* based on plastid (Cameron, 2004a,b; Cameron & Molina, 2006; Pansarin, 2008; Cameron 2009; Bouetard *et al.*, 2010) or mitochondrial loci (Cameron, 2009), or equally related to *Pogonia–Isotria* based on nrITS (Cameron, 2009; Pansarin, 2016). Previous assessments of *Cleistesiopsis* relationships relied on a limited number of markers, whereas our plastid data, based on dozens of loci, support a novel hypothesis placing it as sister to *Isotria*. However, given the incongruence with the nuclear nrITS and mitochondrial topologies and the exceptionally short branches in the Cleistesiopsis + Isotria lineage-a pattern often associated with ILS or hybridization (Mendoza *et al.*, 2020)-this relationship remains the single unresolved case within Pogonieae and clearly warrants further inquiry.

Most cases of genus-level incongruence between our Nuclear and Plastome datasets occur within the Vanilleae tribe and involve its deepest nodes, particularly *Epistephium*, *Clematepistephium*, and *Lecanorchis*, showing substantial divergence from previous inferences. Our results indicate that *Clematepistephium* and *Epistephium* form a highly supported clade despite high gene tree discordance, consistent with elevated levels of ILS, which likely explains the incongruent placements observed in prior studies (e.g., Cameron, 2009). Similarly, the placement of *Lecanorchis* has been historically challenging, varying across nuclear, mitochondrial, and plastid datasets (Cameron, 2009; Givnish *et al.*, 2016; Pansarin, 2016; Giraldo, 2017; Cascales 2023), a pattern we also observed in our nuclear and plastid data. In our Nuclear_Astral analysis, both *Lecanorchis* samples form a clade sister to the holo-mycoheterotrophic vanilloids (*Erythrochis*, *Galeola*, and *Pseudovanilla*) and *Vanilla*, with moderate support but high phylogenomic incongruence. *PhyTop* analysis detected two of five rejections from a non-ILS pattern involving the crown and stem *Lecanorchis* nodes, with moderate-to-high introgression signals. This incongruence appears to stem from deep reticulations rather than ILS, as suggested by non-random quartet distributions and confirmed in our *PhyloNet* analyses, which detected two hybridization events; one between *Galeola* and *Lecanorchis* and another indicating an ancient hybrid origin involving the ancestor of Vanilleae, *Lecanorchis*, and holo-mycoheterotrophic vanilloids + *Vanilla* (Fig. S12). These findings are in accordance with previous suggestions that secondary gene flow has influenced *Lecanorchis* evolution, potentially at different temporal scales (Hwang *et al.*, 2022). Paralogy may also contribute to the phylogenomic discordance. Both *Lecanorchis* accessions exhibited the largest numbers of paralogs during filtering and when paralogs were retained, the samples were recovered in distinct clades, one sister to the holo-mycoheterotrophic *Vanilla* lineages and the other close to *Clematepistephium*. Similar patterns have been reported in lineages affected by whole-genome duplications (Morales-Briones *et al.*, 2022), which may partially explain the historically difficult placement of *Lecanorchis*, particularly as sequence-capture methods using short reads may fail to fully resolve paralogous copies compared with whole-genome sequencing (Rothfels, 2021).

In terms of biogeographic history, our results broadly corroborate previous studies of vanilloid diversification. The DIVALIKE+J reconstruction places the crown node of Pogonieae in the Neotropics at approximately 59.9 Mya (95% HPD: 57.9–61.8), during the Paleocene (Fig. S17, S18), consistent with estimates reported by Givnish *et al*. (2016). The DIVALIKE-J reconstruction places the crown node of Pogonieae in the Neotropics at approximately 59.9 Mya (95% HPD: 57.9–61.8), during the Paleocene (Fig. S19), consistent with estimates reported by Givnish *et al*. (2016). The internal relationships inferred here are also congruent with that study, as well as with Bouetard *et al*. (2010), upon which the former relied for reconstructing major patterns within Vanilloideae. Our reconstruction further supports the hypothesis of an ancient dispersal route from the Neotropics to the Palearctic via the Nearctic. Evidence for this pathway appears in our analysis as a stepwise transition in which the MRCA of Pogonieae originated in the Neotropics, followed by a subsequent range expansion into the Nearctic around 43 Mya (95% HPD: 41.9–45.7). This was followed by an additional expansion yielding a Nearctic–Palearctic ancestral distribution that later underwent vicariance, giving rise to the North American *Cleistes* Rich. Ex Lindl.*, Cleistesiopsis*, and *Isotria*, and the Asian *Pogonia*.

This Neartic–Palearctic disjunction is perhaps the most intensively examined biogeographic pattern within Vanilloideae. Most hypotheses invoke intermittent land connections across Beringia and the North Atlantic during the Paleocene (ca. 66 Mya; Cameron & Chase, 1999; Givnish *et al.*, 2010), an interpretation with which our results are also compatible and which aligns with patterns observed in several other plant lineages exhibiting analogous intercontinental disjunctions (e.g. Wen *et al.*, 2016). By contrast, early dispersal from South America into North America has remained more contentious. The GAARlandia hypothesis has historically been invoked to account for northward movement via the Antilles, proposing that colonization was facilitated by a quasi-continuous land bridge or island chain connecting northern South America to the Greater Antilles for 1–2 Myr at the Eocene–Oligocene boundary (Iturralde-Vinent & MacPhee, 1999). However, empirical support for this connection has been lacking (Nieto-Blasquex *et al.*, 2017). More recent tectonic evidence indicates that between 25–45 Mya the Caribbean region experienced substantial uplift, forming a transient land corridor linking South America with the proto-Caribbean realm (Montheli *et al.*, 2025). This interval is congruent with our inferred timing for a shared ancestral distribution spanning South and North America, suggesting that this tectonic corridor may have served as a conduit for early Pogonieae migration. Interestingly, no extant vanilloid lineages other than *Vanilla* occur in the Antilles. This absence may imply that ancestral Pogonieae traversing a South America–Caribbean–North America route subsequently went extinct in the Caribbean region, leaving no surviving representatives of that dispersal pathway.

Our vanilloid topology departs from earlier reconstructions in several key clades, and these differences, in turn, yield novel biogeographic insights. Previous studies recovered two independent LDD events involving non-*Vanilla* vanilloids from the Neotropics to Australasia and to the Indo-Malayan region (Givnish *et al.*, 2016). In contrast, our analysis identified three LDD events, all occurring near the Paleocene boundary (ca. 60 Mya) (Fig. S17, S18). These comprise: (1) a Neotropical–Australasian dispersal giving rise to *Epistephium* and *Clematepistephium*; (2) a second dispersal from the Neotropics to an Australasia–Indo-Malayan ancestral range within the holo-mycoheterotrophic vanilloids (*Cyrtosia*, *Erythrorchis*, and *Pseudovanilla*); and (3) a third event from the Neotropics to a Palearctic–Indo-Malayan ancestral distribution in *Lecanorchis*. This latter event has not been proposed in previous biogeographic assessments. As noted above, our inferred placement of *Lecanorchis* departs substantially from prior studies. Contrary to earlier reconstructions that consistently embedded *Lecanorchis* within the holo-mycoheterotrophic vanilloid clade (Cameron, 2009; Givnish *et al.*, 2016; Zhou *et al.*, 2023) our analyses place it as sister to the remainder of Vanilleae (excluding *Clematepistephium* and *Epistephium*). Remarkably, this topology is more congruent with a recent phylogenomic study employing the Angiosperms353 target-capture dataset (Pérez-Escobar *et al.*, 2024), which also recovered *Lecanorchis* at a deeper position within Vanilloideae. The concordance between our Astral inference—robust to ILS—and patterns recovered in genome-scale analyses suggests that the nuclear genomic signal captured here may better represent the evolutionary history of *Lecanorchis* than earlier plastid-dominated datasets. Nonetheless, we interpret this result with caution. Additional sampling within *Lecanorchis* and expanded genomic representation will be essential for corroborating the extra LDD event inferred here and for resolving the evolutionary trajectory of this enigmatic lineage.

## Conclusions

This study presents the first comprehensive phylogenomic framework for *Vanilla*. By integrating extensive taxon sampling with nuclear and plastid data, we resolve longstanding phylogenetic uncertainties, clarify major clades, and resolve species-level relationships, confirming some previous hypotheses while revealing novel ones. In particular, we clarify the placement of the widely cultivated *V. planifolia* and, for the first time, establish the position of the monospecific *Vanilla* subg. *Gondwana*, endemic to French Guiana and closely related to predominantly Old-World *Vanilla*. Our analyses further indicate that cyto-nuclear and gene–species tree incongruences in *Vanilla* are primarily driven by ILS, which is particularly influential within the vanillin-producing clade (*Vanilla* subg. *Vanilla*). We also found evidence of both ancient and recent hybridization events, including a natural hybrid endemic to the Yucatán Peninsula.

Our dating and biogeographic analyses support an origin for *Vanilla* in the Guiana Shield during the Oligocene (∼30 Mya), with the region acting as a major center of diversification. While corroborating previous long-distance dispersal events, we provide new insights into Neotropical biogeographical dynamics, showing that most diversification and dispersal occurred within the last 5 My and that Andean uplift facilitated, rather than impeded, dispersal between Amazonia, the Pacific lowlands, and Central America.

We also offer novel insights into the relationships of *Vanilla* allies, particularly clarifying previously challenging genera such as *Clemaepistephium* and *Lecanorchis*, and proposing a new biogeographical history for the entire Vanilloideae subfamily, including an additional long-distance dispersal event not suggested in earlier studies. Collectively, these findings establish a robust evolutionary framework for *Vanilla*, providing a foundation for future taxonomic revisions, comparative trait analyses, and a deeper understanding of the historical processes shaping one of the most economically and biologically significant orchid genera.

## Acknowledgments

We are particularly grateful to the Department of Botany at the University of Wisconsin–Madison for its financial support, which enabled visits to the AMO, F, MEXU, MO, NY, and US herbaria, where we gathered and curated material for DNA sampling. We also extend our appreciation to the Graduate Student Research Award Committee and, in particular, to Mr.

Theodore S. Cochrane at UW–Madison for their generous assistance. The first author further acknowledges funding provided by the American Society of Plant Taxonomists (ASPT), the Botanical Society of America (BSA), the Systematics Association, and the American Orchid Society (AOS). Additional support for the first author’s research was furnished by the Consejo Nacional de Ciencia, Tecnología e Innovación (CONCYTEC)–PROCIENCIA through the program “E041-2023-01 Basic Research Projects” (contract number PE501082530-2023). AD also thanks Inkaterra Asociación for logistical support throughout this research, especially José Koechlin, José Purisaca, Elizabeth Hilario, Nayli Pizarro, Ruth Escobedo, and Manuel Bisso. We are likewise sincerely grateful to the many colleagues who generously shared *Vanilla* material, with special appreciation to Jason Wells, Eduardo Vilcatoma, Joyser Pizango, Arturo Rivas, Sergio Olortegui, Moisés Ródenas, and Manuel Huinga. Finally, we would like to thank to the different Biodiversity agencies that granted collection permits for our work, specially to the Sistema Nacional de Gestao do Patrimonio Genetico e do Conhecimiento tradicional Associado (permits SISGEN-A63D1C7, R-496EF7), Secretaria de Ambiente, Alcadia Mayor de Bogota, Colombia (CITES 47988), University of Costa Rica/Jardin Botanico Lankester (CITES non commercial material transfer agreement), Servicio Nacional Forestal y de Fauna Silvestre, Perú (N° 000144-2023-MIDAGRI-SERFOR-DGGSPFFS-DGSPF) and Ministerio del Ambiente, Agua y Transicion Ecologica, Ecuador (CITES 23EC0000108E).

## Authors contributions

AD-P, OP-E, were involved in conceptualization. AD-P, OP-E, and APK were involved in methodology and formal analysis. AD-P and NM-R were involved in data curation. AD-P, OP-E, APK, AA, JPJ, NM-R, OJF, AB, XW, MEE, MRM, WD, GC, GG, EH, ML-R, YR, LV, HG, LB, GI, AP, MJ, SO, and KK were involved in resources. OAP-E, AD-P, and KC were involved in software. AD-P, OP-E, APK, AA and NM-R were involved in investigation. AD-P, NM-R, OP-E, AA were involved in funding acquisition. AD-P, NM-R, and OP-E were involved in visualization. AD-P, NM-R, were involved in writing – original draft. AD-P, OP-E, APK, AA, JPJ, NM-R, OJF, AB, XW, and MEE were involved in writing – review and editing. AD-P, and OP-E are lead authors on this work. AD-P, OP-E, AA and KC are senior authors on this work.

## Data availability

The data supporting the findings of this study will be openly available in the NCBI Sequence Read Archive (https://www.ncbi.nlm.nih.gov/) under BioProject number TBD. Alignments for Angiosperms353, tree files, and gene and species trees, along with metadata associated with the biogeographical analyses, will be provided via a FigShare link (TBD).

